# Engineering the citrus phyllosphere microbiome for enhanced disease resistance to bacterial canker

**DOI:** 10.1101/2025.11.25.690529

**Authors:** Jinyun Li, Jin Xu, Donghwan Lee, Connor Hendrich, Zhiqian Pang, Wenting Wang, Ma Andrea Cruz Gatmaitan, Yu Feng, Javier L. Dalmendray, Yucheng Duan, Manyu Zhang, Milica Calovic, Chris Oswalt, Nian Wang

## Abstract

Engineering phyllosphere microbiomes toward plant protection against pathogens on crops has shown promising effects. Here we tested the effect of application of beneficial bacteria individually and as a synthetic community (SynCom) on the phyllosphere microbiome and management of bacterial canker caused by *Xanthomonas citri* subsp. *citri* (Xcc). Foliar spray of a SynCom consisting of three citrus endophytic bacterial strains (*Paenibacillus* sp. ATY16, *Bacillus megaterium* PT6, and *Bacillus subtilis* PT26A, SynCom1) with mechanistically complemented plant-beneficial properties exerted a more profound effect on the leaf microbiota and suppression of citrus canker than single strains. Cultivation-dependent and bacterial 16S rRNA gene-sequence profiling analyses revealed that leaf endophytic *Bradyrhizobium, Brevibacillus, Cellvibrio*, *Flavobacterium,* and *Pseudomonas* were significantly enriched in the SynCom treated plants, in addition to *Bacillus* and *Paenibacillus*. The enriched *Brevibacillus* and *Pseudomonas* spp. were isolated, whole-genome sequenced, and confirmed to possess multiple beneficial traits including antimicrobial activity against Xcc, Moreover, the expression of citrus defense genes was induced by the SynCom1 inoculation. Foliar application of SynCom1 and a newly assembled SynCom (SynCom2; SynCom1 with representative *Brevibacillus* and *Pseudomonas* isolates) prior to Xcc inoculation significantly reduced the citrus canker disease severity in greenhouse assays, and SynCom2 had a better efficacy, comparable to a copper bactericide. Three-year field tests showed that both SynCom1 and SynCom2 effectively controlled citrus canker, with a 50% reduction of foliar and fruit canker incidence. Collectively, these findings provide novel insights and valuable clues for microbiome engineering to serve as a sustainable strategy for the control of phyllosphere pathogens on crops.

## Introduction

Plants are colonized by a diverse microbiota, with some microorganisms residing the above ground parts (phyllosphere) and others associating with below ground parts, i.e., rhizosphere and root. Rhizosphere and root-colonizing microbial communities have been extensively studied, and their assembly is shaped by a variety of factors, including plant host genetic variations (Oyserman et al. 2022; Rodríguez et al. 2025; Xu et al. 2023), environmental changes (Cook et al., 2023; Flemer et al., 2022; Wang et al., 2020), and pathogen infection (Berendsen et al. 2018; Flemer et al. 2022; Gao et al. 2021; Trivedi et al. 2012). Rhizosphere and root-colonizing microbes provide host plants with various benefits, such as enhancing disease resistance, stress tolerance, and growth promotion (Castrillo et al. 2017; Kwak et al. 2018; Trivedi et al. 2020). Manipulating the rhizosphere microbiome through microbial amendment and other management practices to improve plant health and productivity and sustainability has been increasingly evaluated and adopted in agricultural and forest production systems (Beattie et al., 2024; Trivedi et al. 2020). Many studies have explored the processes that are responsible for determining the compositions of phyllosphere microbiomes and how they contribute to plant health and fitness (Beattie et al., 2024; Ehau-Taumaunu and Hockett, 2023; Liu et al. 2020; Lombardo et al. 2025; Zhan et al., 2022).

The phylloshpere microbiome consists of epiphytes (microorganisms living on the surface of plants) and endophytes (residing inside the plants) and, plays a pivotal role in maintaining plant health. It is well known that the phylloshpere microbiome may suppress plant diseases through antimicrobial activity against the pathogens, competition for space and resources, and activation of plant immune defense (Alizadeh 2025; Sohrabi et al. 2023). Moreover, it was reported that the homeostasis of endophytic phyllosphere bacterial communities is critical to plant health (Chen et al. 2020). Arabidopsis plants that are defective in key components of innate immunity pathways harbor restructured endophytic phyllosphere microbiomes, leading to leaf-tissue damage associated with dysbiosis (Chen et al. 2020). Interestingly, increasing evidence has shown that certain shifts of microbiota homeostasis in plants under biotic stresses can promote disease resistance (Paasch and He 2021). For example, infection of Arabidopsis leaves by *Hyaloperonospora arabidopsidis* causing downy mildew resulted in the enrichment of specific rhizosphere bacteria that could activate plant resistance against the disease (Berendsen et al. 2018). Additionally, selectively transferring phyllosphere microbiomes from field-grown tomato to greenhouse tomato plants and successive passages of the microbial community including the pathogen *Pseudomonas syringae* pathovar *tomato* between tomato plants could suppress the bacterial speck disease on tomato (Ehau-Taumaunu and Hockett, 2023). Further investigations suggested that this effectiveness of disease suppression was potentially due to the development of disease-suppressive microbial communities resulted from changes in specific microbial populations, e.g., the selection and enrichment of antagonistic populations during the prolonged interaction between the microbial community members and the pathogen (Ehau-Taumaunu et al. 2025). It also has been shown that synthetic communities (SynComs) consisting of members of the core phyllosphere microbiota can protect *Arabidopsis thaliana* against the foliar bacterial pathogen *Pseudomonas syringae* pv. *tomato* DC3000 in a gnotobiotic system (Vogel et al. 2021). Collectively, these studies revealed the great potential of engineering the phyllosphere microbiome for protecting plants against foliar pathogens. However, most previous studies have focused on model systems involving *Arabidopsis* under controlled experimental conditions and, investigations of phyllosphere microbiome engineering in crop plants are limited and at an early stage of development (Alizadeh 2025; Hirata et al. 2025; Northen et al. 2024). More research is needed to develop vigorous biocontrol agents associated with phyllosphere microbial communities for effective control of pathogens in crop fields (Sohrabi et al. 2023).

Citrus is one of the top three perennial fruit crops globally (Wu et al. 2018). Citrus canker caused by *Xanthomonas citri* subsp. *citri* (Xcc; syn. *X. axonopodis* pv. *citri*) is a devastating disease of citrus worldwide (Martins et al. 2020). All commercial citrus cultivars are susceptible to citrus canker (Graham et al. 2004; Ference et al. 2018; Patané et al. 2019). Xcc enters the phyllosphere tissues through stomata and wounds and colonizes the intercellular space to trigger disease progression (Graham et al. 2004). Canker infestation significantly alters the composition and structure of the endophytic bacterial and fungal communities (Huang et al. 2023). Citrus canker symptoms typically include necrotic, erumpent lesions on leaves and fruits with raised, brown, water-soaked margins (Graham et al. 2004). Premature fruit drop, twig dieback, and general tree decline often occur on heavily infected trees. Citrus canker causes significant economic losses worldwide, due to the decrease of fruit quality and yield fruits and the costs for disease control (Gottwald et al. 2002; Martins et al. 2020). In addition, citrus canker has a serious impact on commerce resulting from restrictions to international citrus trade from canker-affected areas because Xcc is designated a quarantine pathogen in many countries (Gottwald et al., 2002; Martins et al. 2020). Foliar spray with copper-based bactericides is currently the most adopted control approach for citrus canker, because other promising measures, such as the use of genome-edited, transgene free canker-resistant citrus cultivars are still far from practical use (Huang et al. 2023; Su et al. 2023) and biological control exhibited limited and/or inconsistent efficacy in the field (Daungfu et al. 2019). However, copper resistance in pathogen and non-pathogen microbial populations have emerged (Behlau et al. 2012; Behlau et al. 2013) and concerns on pollution of the environment arise (Alva et al. 1995; Fan et al. 2011; Graham et al. 1986; Lombardo et al. 2026), as is common with other plant bacterial pathogens (Sundin and Wang, 2018). Effective and eco-friendly disease management approaches of citrus canker are urgently needed.

Microbiome engineering, especially ecologically informed microbiome engineering, aims to manipulate and recreate host-associated microbial communities that harbor substantial biodiversity for increased resilience to ensure colonization and persistence in hosts and sustained functionality to improve ecosystem function (Henry and Bergelson, 2025). It has emerged as a promising environmentally and economically sustainable strategy to enhance plant health and productivity. There is growing evidence that manipulation of the plant microbiome through changes in management practices and/or microbial amendment may suppress phytopathogens and/or promote host-beneficial traits (Alizadeh 2025; Bazany et al. 2022; Cui et al. 2021; Kaur et al. 2022; Tao et al. 2023). In particular, the impacts of phyllosphere microbiomes on plant health and exploiting their functional roles and associated mechanisms in disease suppression have gained increasing attention in the scientific community in the past several years (Berg and Koskella 2018; Chen et al 2020; Ehau-Taumaunu and Hockett 2023; Ehau-Taumaunu et al. 2025; Li et al. 2022; Lombardo et al. 2025; Zhan et al. 2022). Recent studies showed that engineering phyllosphere microbiomes could effectively control plant diseases, such as the fire blight disease in apples (Cui et al. 2021), rice blast (Thapa et al. 2021), and bacterial speck disease in tomato (Berg and Koskella 2018; Ehau-Taumaunu and Hockett 2023).

In this study, we hypothesized that manipulating the phyllosphere microbiome can provide citrus plants with protection against Xcc and suppress citrus canker. We constructed a SynCom (designated as SynCom1) consisting of citrus endophytic bacteria possessing multiple mechanistically complemented plant-beneficial properties, including producing antimicrobials, the plant hormone auxin (IAA: indole-3-acetic acid) and siderophore, and solubilizing phosphate (P). We tested its ability to reshape the citrus phyllosphere microbiome, using cultivation- and bacterial 16S ribosomal RNA (rRNA) amplicon profiling-based approaches. Interestingly, application of SynCom1 recruited other beneficial bacteria, which we isolated and utilized to construct another SynCom (designated as SynCom2). We then evaluated both SynComs for suppression of citrus canker under greenhouse and field conditions to expand our understanding of whether and how microbiome engineering achieves effective control of phyllosphere pathogens.

## Materials and Methods

### Bacterial strains and growth conditions

The citrus endophytic bacteria *Paenibacillus* sp. ATY16, *Bacillus megaterium* PT6, and *Bacillus subtilis* PT26A were used for the construction of SynCom1. The three bacteria were isolated from the root of healthy-looking citrus trees in citrus groves from Florida, USA and evaluated for plant-beneficial properties (summarized in **Supplementary Table 1**) (Trivedi et al., 2011; Li et al. 2013). ATY 16 inhibits Xcc growth *in vitro*. All three strains can produce auxin (indole-3-acetic acid, IAA) and siderophore and solubilize phosphate (P). Relatively, PT26A produced the highest levels of IAA and siderophore, and PT6 had best performance in P solubilization. These three bacteria were compatible with each other as determined by co-culture assays (Li et al. 2013). The whole genomes of strain ATY16, PT6, and PT26A were sequenced and deposited in the NCBI database under the accession number PRJNA392438, PRJNA392439, and PRJNA392442, respectively. Xcc strain 306 causing bacterial canker in citrus (Rybak et al., 2009) was used for inoculation in greenhouse assays. All bacterial strains were stored at –80°C in 30% glycerol, and were routinely grown in nutrient broth (NB) (BD, Sparks, MD, USA) or nutrient agar (NA) medium at 28°C.

### Plant materials and growth conditions

One-year old Valencia sweet orange (*Citrus sinensis*) plants were grown in Metro-Mix soil (Sun Gro Horticulture, Agawam, MA, USA) in 3.8 L pots and maintained in a greenhouse at a temperature of 20 to 30°C and 50 to 60% relative humidity. Plants were watered every other day and fertilized every two weeks with Peters Professional 20-10-20 (N:20%, P:20% and K: 20%) (The Scotts Company, Marysville, Ohio) at a rate of 0.5g/L in water. Five weeks prior to treatment applications, plants were cut back to approximately 30 cm, and shoots were allowed to grow to 20-30 cm to obtain 100% expanded immature leaves suitable for bacterial inoculation through foliar spray.

### Plant inoculation with SynCom1 and single strain

Bacterial strains of ATY16, PT6, and PT26A were cultured in NB medium (for ATY16, adding 0.5% yeast extract and 0.5% xylan to NB) at 28°C with shaking at 180 rpm overnight. The culture was adjusted with sterilized deionized water (SDW) to an optical density at 600 nm (OD_600_) = 0.5 (approximately 10^9^ colony-forming units (CFU)/ml). The bacterial inoculum, regardless of SynCom1or single strain, contained equal amount of the total bacteria (3 × 10^8^ CFU/ml). For individual bacterium, overnight culture (300 ml, adjusting to OD_600_ = 0.5) was directly diluted into 1.0 liters of SDW and used for plant inoculation. For SynCom1, the three ingredient bacteria were mixed at equal amount right before the application. A total of 20 Valencia sweet orange plants in greenhouse were randomly arranged into five treatment groups, with 4 replicate plants in each group. The treatments were: (i) ATY16, (ii) PT6, (iii) PT26A, (iv) SynCom1 (i.e., ATY16 + PT6 + PT26A), and (v) SDW, which was used as an untreated control. The treatments were applied once when at least 12 fully expanded immature leaves were present in each plant. Approximately 200 ml of bacterial suspension at the final concentration of 3 × 10^8^ CFU/ml was applied to each plant with a handheld mist spray bottle until runoff so that all leaves received the inoculum. After treatments, the plants were maintained in a greenhouse at a temperature of 20 to 30°C and 50-60% relative humidity.

### Leaf sampling for quantification of culturable bacteria, isolation of endophytic bacteria and 16S rRNA gene-sequence profiling

Sampling was conducted 2 and 14 days after bacterial inoculation. At each timepoint, six leaves were randomly collected from each plant with sterilized scissors and tweezers and placed in a sterile zipper bag (Whirl-Pak, Madison, Wisconsin, USA). The samples were kept on ice during transportation and stored at 4°C until processing for the experiments, which were completed within 2 days after sampling.

#### Quantification of culturable bacteria

To quantify culturable total bacterial communities, two leaves from each treatment were weighed and ground in 5 ml sterile 10 mM MgCl_2_ using sterile mortar and pestle. Serial 10-fold dilutions were made with homogenized samples and 100 μl of the 10^-3^ to 10^-6^ dilutions were plated on NA plates. Bacterial colonies were counted 5 days after the NA plates were incubated at room temperature. CFUs were normalized to fresh tissue weight. Endophytic leaf bacterial communities were determined following the same protocol except that leaves were surface-sterilized in 75% ethanol for 1 min and rinsed in SDW three times and air-dried to evaporate surface water. Serial dilutions were made and 100 μl of the 10^−2^ to 10^−5^ dilutions were plated on NA plates.

#### Isolation and identification of leaf endophytic bacteria

To isolate endophytic bacteria, leaf samples were processed as described above. For the isolation of bacteria belonging to *Bacillus,* bacterial suspensions were treated at 60°C for 15 min, then 10-fold diluted to 10^-5^ and 100 μl of the 10^-2^ to 10^-5^ dilutions were plated NA plates and incubated at 28°C for 2 days. For isolation of bacteria of *Pseudomonas*, bacterial suspensions were diluted to 10^-5^ and 100 μl of the 10^-2^ to 10^-5^ dilutions were plated on King’s B (KB) (Sigma) agar plates. The plates were incubated at 28°C for 3 days. Bacterial isolates were obtained from agar plates with between 50 and 100 colonies by picking colonies differentiated by their morphology. All the isolates were purified twice by streaking onto fresh NA agar plates and then stored at −80°C in medium broth containing 30% of glycerol for further use. Bacterial isolates of interest were identified based on the full-length 16S rRNA gene sequence obtained through polymerase chain reaction (PCR) using the 27F/1492R primer pair and subsequent amplicon sequencing analysis, as described by Trivedi et al. (2011).

#### DNA extraction for 16S rRNA gene-sequence profiling

For the analysis of endophytic bacterial communities, leaf samples were first sonicated in water bath for 5 min to remove epiphytic microbes, followed by surface-sterilization with 5% (v/v) bleach for 2 min and rinsing with SDW three times. After air-drying to remove surface water, six leaf discs (0.60-cm in diameter) were cut from a leaf using a sterile cork borer. A total of 12 leaf discs from two leaves/plant were collected in a 2 mL screw cap tube that contained two sterile 4.5-mm stainless steel beads as one biological replicate. Then the samples were ground in liquid nitrogen using the Tissuelyser II (QIAGEN). To prepare DNA for bacterial 16S rRNA gene-based community analysis, total DNA from 0.1g pulverized leaf tissues was extracted using DNeasy PowerSoil Pro Kit (Qiagen), following the manufacturer’s instructions. The amount and purity of isolated DNA samples were determined using the Synergy multi-mode reader (BioTek Instruments Inc. Winooski, VT).

### 16S rRNA gene amplicon sequencing and data analysis

Isolated DNA was used as a template to PCR amplify the V5-V7 region of the bacterial 16S rRNA gene using the 799F/1193R primer set (799F: AACMGGATTAGATACCCKG and 1193R: ACGTCATCCCCACCTTCC) (Beckers et al. 2016) for 16S rDNA amplicon library preparation and sequencing per the manufacturer’s protocol using the Illumina NovaSeq6000 platform at Novogene, Sacramento, CA. After quality control, quantification, and normalization of the DNA libraries, 250-bp paired-end (PE) reads were produced from the NovaSeq6000 platform according to the manufacturer’s instructions.

The raw reads from amplicon sequencing were used to generate high quality reads by removing adaptor sequences, trimming, and removing low quality reads (reads with N bases and a minimum quality threshold of 20). Then DADA2 program (Callahan et al., 2016) was used to filter and denoise sequences, remove chimaeras, identify representative sequences of amplicon sequence variants (ASVs). To obtain the taxonomic information of the ASVs, representative sequences of each ASV were generated and aligned against the SILVA databases (Quast et al., 2013), using the Ribosomal Database Project (RDP) Classifier (Wang et al., 2007; Wang et al. 2024). The ASVs defined as unknown, chloroplast, mitochondria, or plants, were removed. The relative abundance for taxa (ASVs and phylum) were computed based on the read counts for each taxon across samples using the Total-Sum Scaling (TSS) method. The alpha diversities of leaf microbiomes were estimated based on Shannon’s index calculated with relative abundance of ASV in individual bacterial communities using the rarefaction method. To present the dissimilarities of bacterial communities among different samples (beta-diversity), the weighted UniFrac distance was calculated and plotted using principal coordinate analysis (PCoA). Based on the read abundance profiles, ASVs with significantly differential abundances between treated and untreated samples were determined using DESeq2 with a negative binomial generalized linear model. The read count matrix for DESeq2 analysis was normalized using the DESeqVS method (Weiss et al. 2017). The co-occurrence network analysis was performed using the identified ASVs and the non-parametric Spearman correlation analysis. The co-occurrence correlations were regarded as robust only if the Spearman’s correlation coefficient (ρ) <−0.70 or >0.70 and significant *P* value < 0.05 and the robust correlations were used to construct the network. The network was visualized using the ggnet2 package in R program.

### Assays for expression of plant defense-related genes in response to bacterial treatment

Bacterial treatments and foliar spray application were performed as described for leaf microbiome analysis. Citrus leaves for RNA extraction were collected at 0 (immediately prior to bacterial treatment), 4, 10, 24, 48, and 72 hours after the bacterial treatments. At each sampling time point, two leaves were randomly collected from each treated plant and pooled as one biological replicate and immediately placed into liquid nitrogen. The samples were kept in liquid nitrogen during transportation and stored at −80°C until RNA isolation. Plant RNA extraction and relative expression of seven defense-related genes involved in salicylic acid (SA)- and jasmonic acid (JA)-dependent signaling pathways (**Supplementary Table 2**) using reverse transcription quantitative PCR (RT-qPCR) analysis were conducted as previously described (Riera et al., 2018), with the housekeeping gene encoding a glyceraldehyde 3-phosphate dehydrogenase (GAPDH) as an endogenous control. The relative fold change of target gene expression in bacteria treated plants was calculated and normalized against its expression levels prior to bacterial inoculation (0 h after the bacterial treatments) using the ΔΔCt method as previously described (Livak and Schmittgen 2001). SDW was used as mock treatment. The experiments were repeated twice with four biological replicates.

### In vitro testing of bacterial isolates for antibacterial activity against *X. citri* subsp. *citri* and plant growth promoting activities

The antibacterial activity of individual leaf endophytic bacterial isolates against Xcc strain 306 in vitro were tested by an antagonistic assay using dual culture technique. Briefly, 200 µl of Xcc strain 306 solution (OD_600nm_ = 0.3, approximately 5 × 10^8^ CFU/mL) was spread on the NA plate. Three 5 µl droplets of individual tested bacteria (OD_600nm_ = 0.3) were placed in one plate as replicates. Plates were incubated at 28°C for 48 h. Positive antibacterial activity was recorded when an inhibitory halo was present around the antagonistic bacterium. Assays for indoleacetic acid (IAA) and siderophore production by bacteria tested were conducted as previously described (Trivedi et al., 2011). The experiments were repeated three times.

### Bacterial DNA isolation, genome sequencing, assembly, and annotation

Bacterial isolates (EBL1 and EPL5) were grown overnight in 5 mL of NB medium at 28°C with 180 rpm agitation. The cultures were centrifuged at 13,000 g for 5 min and DNA was extracted using the Wizard Genomic DNA Purification Kit (Promega) following the manufacturer’s instructions. DNA quality and quantity were measured using Agilent 2100 BioAnalyzer 220 (Agilent Technologies, Inc., Santa Clara, CA, USA). The whole genome sequencing of the two strains was performed using an Illumina Hiseq 2500 platform with a paired-end 150-bp sequencing strategy (Illumina, Hayward, CA, USA). All de novo assemblies were performed on CLC Genomics Workbench 6.0 (CLC bio, Cambridge, MA, USA) with default parameters. Genome functional annotation was completed using the Prokka automatic pipeline (Seemann, 2014).

### Identification of bacterial genes involved in antimicrobial production and beneficial traits

Prediction of putative antimicrobial biosynthesis clusters in bacterial genomes was performed using antiSMASH program (version 7.0.1.) (Weber et al. 2015). Protein BLAST analyses were conducted against the Kyoto Encyclopedia of Genes and Genomes (KEGG) Orthology (KO) database using DIAMOND (Buchfink et al. 2015) to obtain information of biological functions and pathways associated with bacterial genes. Specifically, protein families related to phosphate solubilization, nitrogen fixation, production of siderophores, phytohormone production, and synthesis of volatile organic compounds (VOCs) were searched in all the bacterial genomes obtained in this study.

### Biocontrol of citrus canker in greenhouse

The preparation of bacterial culture and inoculum and the foliar spray application were performed as described for leaf microbiome analysis. The bacterial treatments included the single strains ATY16, PT6, PT26A, EBL1 and EPL5, and SynCom1 and SynCom2 (i.e., ATY16 + PT6 + PT26A+ EBL1 + EPL5). Inoculation of Xcc 306 was conducted 2 days after bacterial treatment, i.e., at the same time as the first sampling for leaf microbiomes. Xcc 306 was inoculated by spraying a bacterial suspension (5.0 × 10^8^ CFU/ml) onto the leaf abaxial surfaces as described by Li and Wang (2011). Additional treatments of the bactericide copper hydroxide (Kocide 3000; 30% metallic Cu a.i.; DuPont) (200 ppm) and SDW were included with four replicated plants for each treatment to serve as positive and negative control for suppression of citrus canker, respectively. Copper hydroxide and SDW were applied through foliar spray on the same day 2 h prior to Xcc 306 inoculation. The plants were then enclosed with plastic bags for 24 h to provide a high relative humidity (>90%) and to favor the opening of stomata for development of canker symptoms. The sprayed leaves were photographed and the number of canker lesions on leaves of each plant were counted with a 10x magnifier at 28 days post Xcc 306 inoculation. Xcc 306 populations in leaves were determined using a dilution-plating method as described by Li and Wang (2011). The experiments were repeated two times.

### Biocontrol of citrus canker under field conditions

We tested the control effect of SynCom1 and SynCom2 against citrus canker in a commercial grove. To prepare bio-formulations of SynCom1 and SynCom2, individual strains of ATY16, PT6, PT26A, EBL1 and EPL5 were grown in NB medium (for ATY16, added 0.5% yeast extract and 0.5% xylan) individually with a 10-liter Winpact bench-top fermenter (Major science, Saratoga, CA, U.S.A) with agitation at 70 rpm for 24 h at 28°C. For both SynCom1 and SynCom2, equal amounts of ingredient strains were mixed at approximately 1 ×10^9^ CFU/ml, and a water-soluble sodium salt of lignin (Sigma, St. Louis, MO, USA) was added at 2.0 g/L under sterile conditions as UV protectant as suggested by Schisler et al. (2004).

In 2022, field trials were conducted in a commercial citrus grove of ‘OLL-8’ sweet orange on ‘US-942’ rootstock (a hybrid of Sunki mandarin (*Citrus reticulata*) and Flying Dragon Trifoliate Orange (*Poncirus trifoliata*)) planted in April 2018 at Lake Alfred, Florida. The grove was established on well-drained Astatula fine sand soil (Typic Quartzipsamment) and >90% trees were naturally infected by Xcc. During the experiments, all trees received standard commercial care including regular irrigation, fertilization, and pest management practices except applications of bactericides, which continued throughout the trial. Treatments were arranged in a randomized complete block design (RCBD) with four treatments replicated five times in blocks of five contiguous trees. Treatments applied were (i) SynCom1 and (ii) SynCom2, with a total dose of 1.0 ×10^9^ CFU/ml in the prepared bio-formulations, (iii) copper hydroxide at 0.57 g a.i. per tree per application (Kocide 3000; 30% metallic Cu a.i.; DuPont), and (iv) water-only spray as the untreated control (UTC). Copper hydroxide was mixed with water. Copper hydroxide and bacterial bio-formulations were applied every three weeks as foliar sprays at 2.0 liters/tree with a handgun sprayer at 1,400 kPa of air pressure. Treatments were initiated after the spring flush in March 2022 and continued until the fall flush in October 2022.

In 2023 and 2024, the treatments were repeated in the same grove as the 2022 trial. The treatments were the same as in 2022, with the addition of two treatments of SynCom1 and SynCom2 rotated with copper hydroxide respectively. Treatments started after the spring flush in March and continued until the fall flush in October 2023 and 2024.

Canker disease severity was evaluated as previously described (Graham and Myers, 2013). Briefly, the incidence of canker-diseased leaves and fruits was evaluated with 100 leaves per four foliar flushes and 50 fruits per tree respectively in November and was expressed as the percentage of the total number of leaves or fruits with canker lesions.

### Statistical analysis

All statistical analyses were performed using SAS V9.4 (SAS Institute Inc., Cary, NC). The data were first tested for normality and homogeneity of variance. When normality and/or homogeneity of variance were not satisfied, the data were log_10_(X+1)-transformed prior to analysis. An one-way analysis of variance (ANOVA) was performed to determine any differences among treatments in population sizes of leaf total and endophytic microbiota, alpha diversity (Shannon’s index) of leaf microbiome, citrus canker severity (number of lesions per leaf) and Xcc strain 306 titers *in planta* in greenhouse tests, and citrus canker disease incidence in field experiments. Multiple comparisons were made using Tukey’s Honest Significant Difference (HSD) test with a significance level at α = 0.05. Permutational multivariate analysis of variance (PERMANOVA) was performed to determine the effects of different factors on the leaf bacterial community dissimilarity using the weighted UniFrac distance (beta diversity).

## Data availability

The raw sequencing reads of 16S rDNA amplicon data and nucleotide sequence and assembled genomes of the three bacterial strains *Brevibacillus agri* EBL1, *Pseudomonas chlororaphis* EPL5, and *Pseudomonas fluorescens* EPL8 were deposited in the National Center for Biotechnology Information (NCBI) database under the accession number PRJNA1238003 and PRJNA1150675, respectively.

## Results

### Citrus leaf microbiome is altered by bacterial inoculation

To understand the effect of bacterial inoculation on phyllosphere microbiome assemblage, we investigated the changes in leaf bacterial communities over a period of two weeks following inoculation with *Paenibacillus* sp. ATY16, *B. megaterium* PT6, *B. subtilis* PT26A and SynCom1 (ATY16 + PT6 + PT26A). Firstly, the population sizes of leaf microbiota were determined using cultivation-dependent analyses at 2- and 14-days post inoculation (DPI). Marked differences in the levels of both total leaf bacteria (including both epiphytic and endophytic bacteria) and endophytic bacteria (estimated after surface sterilization to remove epiphytic bacteria) were observed between the bacteria inoculated and water treated control plants (**Supplementary Figure 1**). Both total bacteria and endophytic bacteria counts in the inoculated plants were approximately 100-fold higher than those in control plants. No significant differences in total or endophytic population sizes were observed among bacterial treatments at either time point. Inoculation with single bacterial strains or SynCom1 conferred similar population sizes of total or endophytic bacterial communities (**Supplementary Figure 1**). The results indicated that both epiphytic and endophytic phyllosphere microbiota were influenced by bacterial inoculation with single bacteria or SynCom1. Given that Xcc colonizes the intercellular space, i.e., live and proliferate endophytically, to trigger citrus canker disease, we focused on the changes of endophytic phyllosphere microbiota elicited by the bacterial inoculation.

**Fig. 1.**
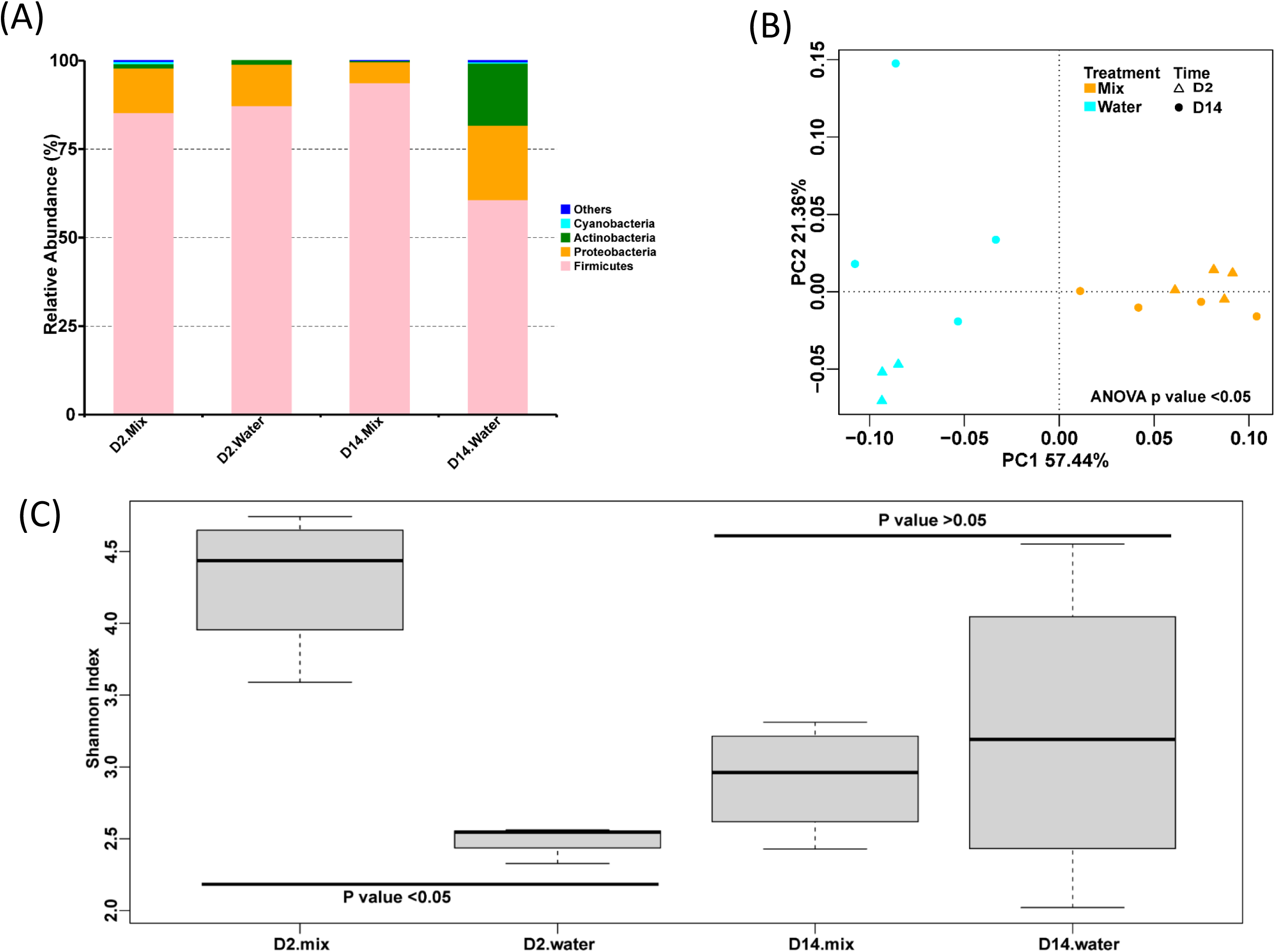
Endophytic leaf microbiomes in the bacterial mixture SynCom1 sprayed and water treated citrus plants. (A) The relative abundance of bacteria at the phylum level obtained from 16S rRNA gene-sequence profiles of endophytic leaf bacteria in citrus plants sprayed with the bacterial mixture SynCom1 or water across time. (B) Principal coordinate analysis (PCoA) based on the weighted UniFrac distance in the bacterial mixture SynCom1 treatment and control samples from each time point. *P*-value was generated by permutational multivariate analysis of variance (PERMANOVA); (C) Shannon indexes (alpha diversity) calculated at the genus level based on from 16S rRNA gene-sequence profiles of endophytic leaf bacteria in citrus plants sprayed with the bacterial mixture SynCom1 or water. In box plots, the center line denotes the median, box edges represent the 75th and 25^th^ percentiles, and whiskers show 1.5× the interquartile range. *P* values were generated using Tukey’s HSD test following one-way ANOVA and *P* < 0.05 was considered as significant different. *n* = 3 biological replicates (water treatment at 2 days post inoculation (DPI)) and *n* = 4 biological replicates (bacterial mixture treatment at 2 and 14 DPI and water treatment at 14 DPI). D2: 2 DPI. Mix: the bacterial mixture SynCom1; D14: 14 DPI. Water: water treatment as untreated control.

Comparative 16S rRNA gene-sequencing analyses were then conducted for citrus leaf endophytic bacterial communities in bacteria inoculated and water treated control plants. In total, 32,673 high-quality 16S rRNA gene reads were obtained and 358 ASVs (average 23 per sample) were identified across treatments and sampling time points. The majority belonged to the phyla Firmicutes (~52 to 94%), Proteobacteria (~6 to 34%), and Actinobacteria (~0.3 to 24%) (**Fig. 1A**; **Supplementary Figures 2A, 3A, and 4A**). Principal coordinate analysis (PCoA) based on weighted UniFrac distance (beta diversity) showed that variations in abundance of the ASVs occurred among samples. The SynCom1 samples clustered together with each other and separated from samples of control, indicating that the community composition of leaf microbiomes between the SynCom1 treatment and control significantly differed at both time points (*P* < 0.05, permutation-based ANOVA) (**Fig. 1B**). Notably, the leaf endophytic bacterial communities in SynCom1 inoculated plants were substantially more diverse than those in water treated control plants at 2 DPI (*P* < 0.05; one-way ANOVA and Tukey’s HSD test), as judged by the Shannon’s index (indicator of alpha diversity) (**Fig. 1C**), and the differences in bacterial community diversity were not observed at 14 DPI. When the plants were treated with single strains, significant differences were detected in the community composition of leaf microbiomes between strain ATY16 or PT26A treatment and control (**Supplementary Figures 2B, 3B, and 4B**). However, there were no significant differences (*P* > 0.05,) in the diversity of leaf endophytic bacterial communities between any single strain treated and control samples at either 2 or 14 DPI (**Supplementary Figures 2C, 3C, and 4C).**

**Fig. 2.**
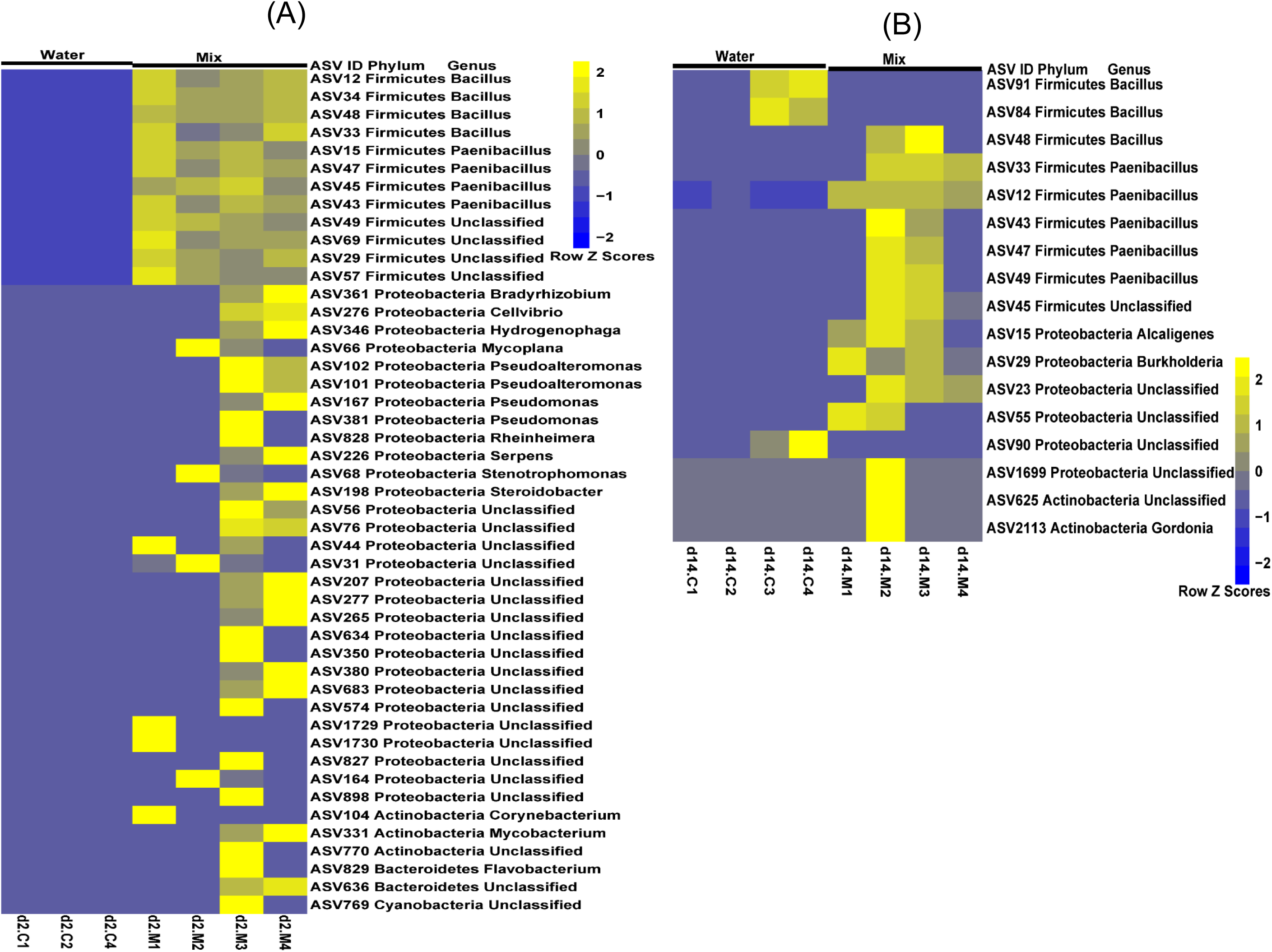
The relative abundance of bacteria at the genus level obtained from 16S rRNA gene-sequence profiles of endophytic leaf bacteria in citrus plants sprayed with the bacterial mixture SynCom1 or water at 2 (A) and 14 (B) days post inoculation (DPI). All the bacteria that showed significantly different abundance (corrected *P* values < 0.05; DESeq2) between the two treatments were presented. Color scale: relative abundance of taxon at row normalization by removing the mean (centering) and dividing by the standard deviation (scaling). The color from blue to yellow represents a relative abundance of each taxon from low to high. *n* = 3 biological replicates (water treatment at 2 days post inoculation (DPI)) and *n* = 4 biological replicates (bacterial mixture treatment at 2 and 14 DPI and water treatment at 14 DPI). d2: 2 DPI. Mix: the bacterial mixture SynCom1; d14: 14 DPI. Water: water treatment as untreated control.

**Fig. 3.**
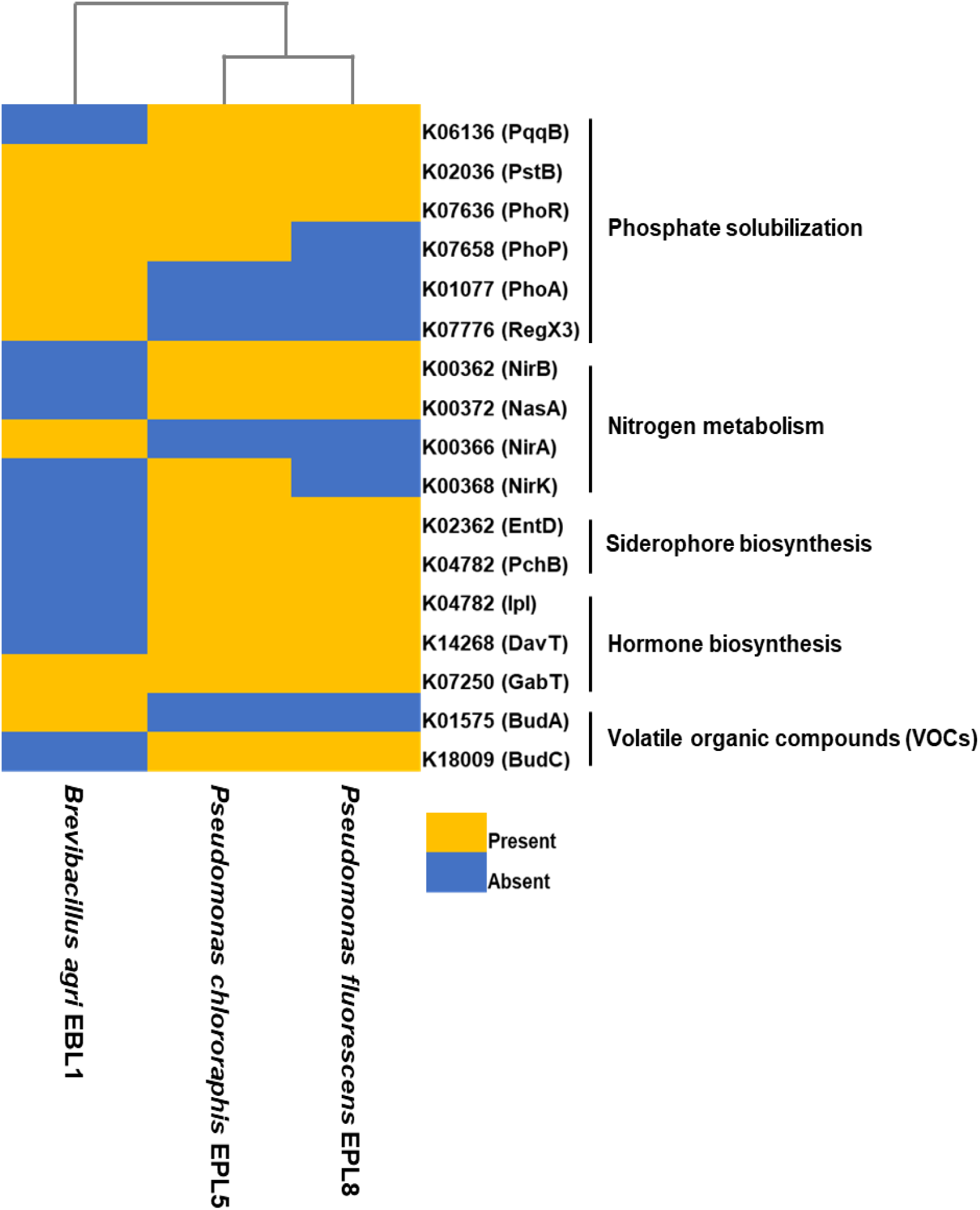
Plant beneficial traits-related genes identified in bacterial isolates. The heat map represents the absent and present of bacterial genes that are putatively involved in phosphate solubilization, nitrogen fixation, siderophore production and iron acquisition, hormone regulation, and production of volatile organic compounds (VOCs).

**Fig. 4.**
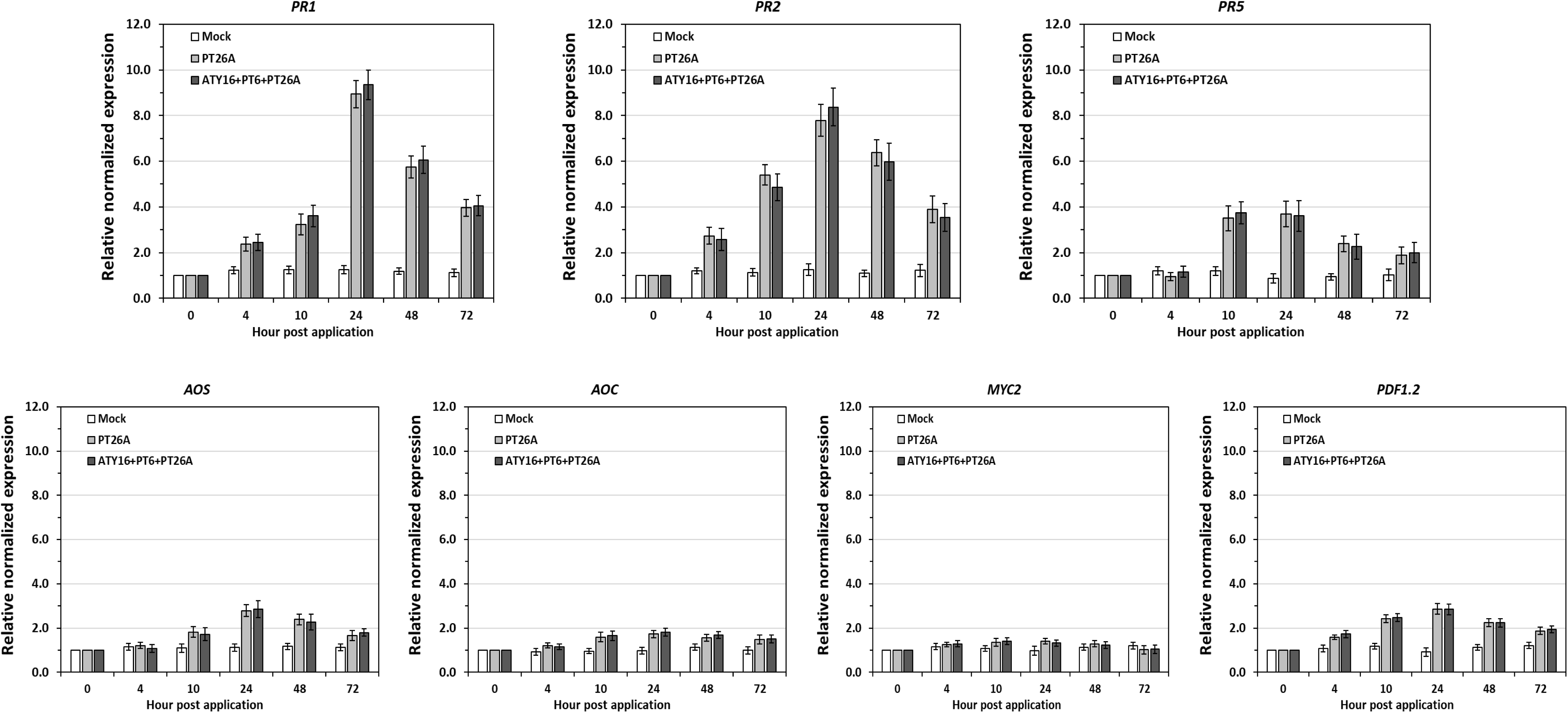
Expression pattern of defense-related genes in leaves of Valencia sweet orange following spray with the indicated bacterial inoculum. Relative transcript abundance of the genes was normalized against their expression levels prior to treatment application (0 hour post application). Error bars indicate standard deviation of three biological replicates. Mock: water. PT26A: *Bacillus subtilis* PT26A. ATY16+PT6+PT26A: the bacterial mixture SynCom1 containing bacteria *Paenibacillus* sp. ATY16, *Bacillus megaterium* PT6, and *Bacillus subtilis* PT26A at equal rations. The experiments were repeated two times with similar results.

Further analysis of endophytic bacterium 16S-profiling data identified significant changes (corrected *P* < 0.05, DESeq2) in specific ASVs between SynCom1 inoculated and control plants. Interestingly, at 2 DPI, there were 47 ASVs substantially more abundant in SynCom1 inoculated plants compared with water treated plants (**Fig. 2A; Supplementary Table 3**). The ASVs belonging to the genera of *Bacillus*, *Paenibacillus*, *Pseudomonas*, *Stenotrophomonas*, *Cellvibrio* and *Bradyrhizobium* were enriched in SynCom1inoculated plants. At 14 DPI, 17 ASVs showed a significantly different relative abundance in SynCom1 inoculated plants compared with control plants (**Fig. 2B; Supplementary Table 4**), although the global bacterial community diversity did not significantly differ between the two groups at this timepoint (**Fig. 1C**). Among them, 14 ASVs belonging to *Bacillus*, *Paenibacillus*, and *Burkholderia* were enriched in SynCom1 inoculated plants (**Fig. 3B**). These observations implied that certain bacteria were enriched in the leaf endophytic microbial communities in response to SynCom1 inoculation. Furthermore, the co-occurrence networks in the SynCom1 treated microbiomes had significantly positive correlation among the ASVs enriched, especially the ASVs from Firmicutes and Proteobacteria, with a higher degree of connection than non-enriched ASVs (**Supplementary Figure 5**). Collectively, these findings indicate that bacterial inoculation significantly affected the citrus leaf microbiome and SynCom1 exhibited a much more profound effect than any single strain tested, resulting in significantly differed leaf bacterial community diversity and structures relative to untreated control.

**Fig. 5.**
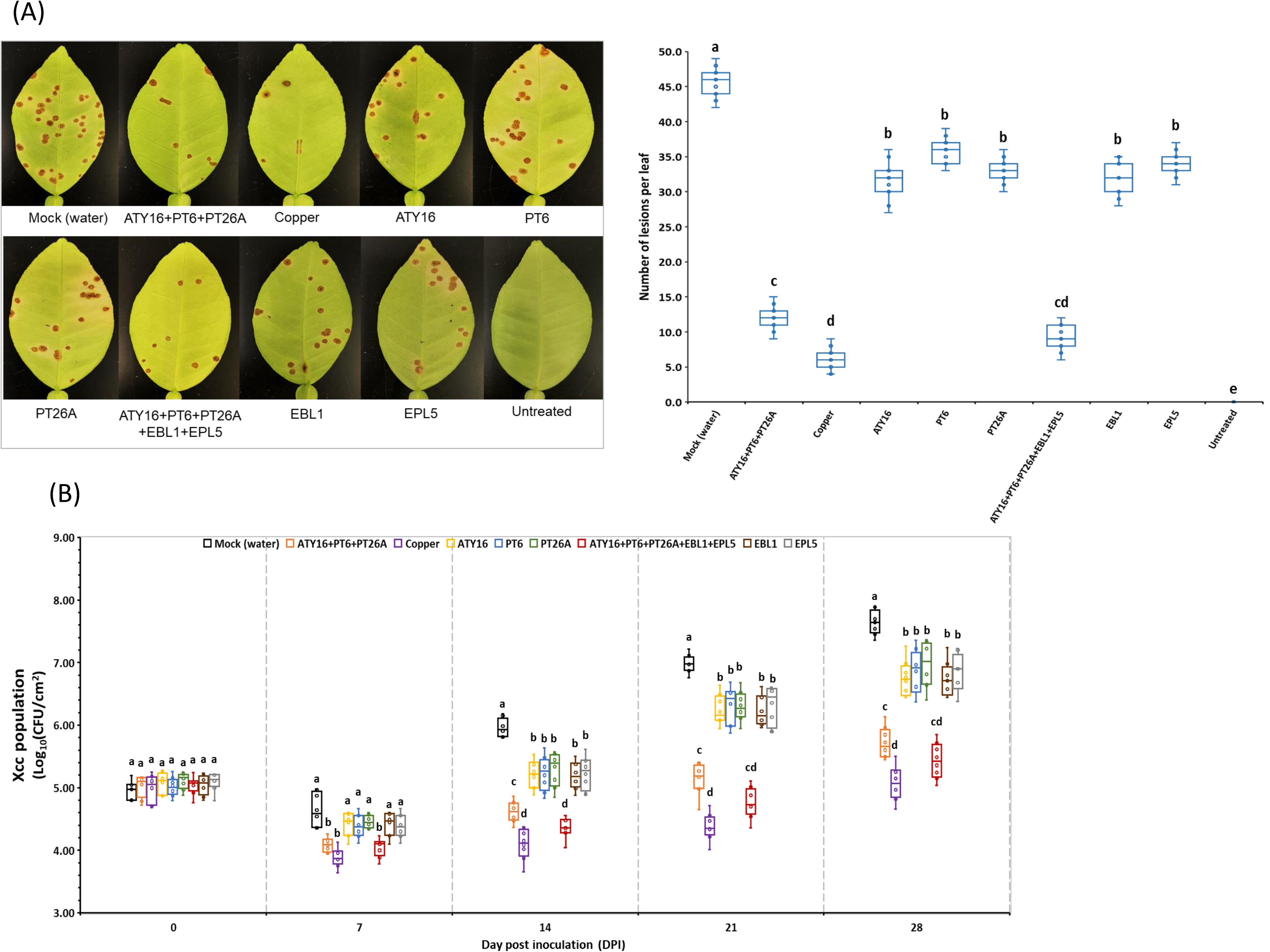
Biocontrol of citrus canker with single bacterial strains and synthetic communities (SynComs). (A) Representative images of leaves showing canker lesions are shown on the left and lesion numbers per leaf under different treatments are shown on the right. Pictures of symptomatic leaves and lesion counts were obtained at 28 days post inoculation (DPI) of *Xanthomonas citri* ssp. *citri* strain 306. Different lowercase letters denote significant differences in lesion counts per leaf among the treatment groups (*P* < 0.05; Tukey’s HSD test at α = 0.05). Combining two independent experiments, *n* = 9 leaves per treatment. (B) Population dynamics of *Xanthomonas citri* ssp. *citri* (Xcc) strain 306. Xcc populations were estimated by homogenizing leaf discs (0.8 cm in diameter) in 0.85% (w/v) NaCl, followed by dilution plating. Data from four independent replicates of each experiment are represented as boxplots with each symbol corresponding to bacterial populations from one citrus leaf. Different letters denote significant differences among the treatments within a single time point (*P* < 0.05; Tukey’s HSD test at α = 0.05). Combining two independent experiments, *n* = 8 leaves per treatment per time point. In box plots, the center line indicates the median, box edges represent the 75^th^ and 25^th^ percentiles, and whiskers show 1.5× the interquartile range.

### Recovery of the enriched leaf microbes in SynCom1 treated citrus plants

To characterize the enriched ASVs belonging to *Bacillus*, *Paenibacillus*, and *Pseudomonas* in SynCom1 treated plants, we isolated the leaf endophytic bacteria. We specifically focused on *Bacillus*, *Paenibacillus*, and *Pseudomonas* since many species of them have beneficial traits for plants. Using broad-spectrum media under selective culture conditions for *Bacillus* and media selective for *Pseudomonas*, 61 isolates were initially obtained according to colony morphology on agar plate. Based on full-length 16S rRNA gene sequences, the 61 isolates were identified to 17 different bacterial species belonging to four genera, including *Bacillus*, *Brevibacillus, Paenibacillus*, and *Pseudomonas* (**Table 1**). Among the 61 isolates, three (EBL4, EBL5, and EBL6) shared 100% 16S rRNA sequence identity to the inoculum *B. subtilis* PT26A, other three (EBL11, EBL12, and EBL13) showed 100% sequence identity to *B. megaterium* PT6, and two (EBL14 and EBL15) had 100% sequence identity to *Paenibacillus* sp. ATY16. The other isolates from the *Bacillus* group were identified to 10 species, including *B. cereus, B. licheniformis, B. ginsengihumi, B. firmus, B. methylotrophicus, B. aryabhattai, B. bombysepticus, Bacillus sp., Brevibacillus agri,* and *Paenibacillus glycanilyticus.* The 16S rRNA gene sequencing analyses also revealed 16 *Pseudomonas* isolates belonging to *P. chlororaphis*, *P. cichorii*, *P. fluorescens*, and *P. putida*. These results indicate that we recovered the three inoculated bacterial species and ASVs belonging to 14 bacterial species that were promoted in the leaves by microbial inoculation.

**Table 1.** Isolates of enriched leaf endophytic bacteria from citrus plants received microbiome manipulation in greenhouse *^Z^*.

| Isolates | Close relative from Genbank (Identity of 16SrRNA) | Antibacterial activity | IAA production | P - solubilization | Siderophore production |
| --- | --- | --- | --- | --- | --- |
| EBL1, 2, 3 | *Brevibacillus agri* (99%) | + | - | + | - |
| EBL4, 5, 6 | *Bacillus subtilis* PT26A (100%) | + | + | + | + |
| EBL7, 8 | *Paenibacillus glycanilyticus* (99%) | + | - | + | - |
| EBL9, 10 | *Bacillus cereus* (98%) | - | + | + | - |
| EBL11, 12, 13 | *Bacillus megaterium* PT6 (100%) | + | + | + | + |
| EBL14, 15 | *Paenibacillus* sp. ATY16 (100%) | + | + | + | + |
| EBL16 | *Bacillus licheniformis* (99%) | - | + | + | + |
| EBL17, 18 | *Bacillus ginsengihumi* (98%) | - | - | + | - |
| EBL19, 20, 21 | *Bacillus methylotrophicus* (98%) | - | - | + | - |
| EBL22, 23, 24, 25 | *Bacillus firmus* (98%) | - | + | + | - |
| EBL26, 27, 28, 29 | *Bacillus aryabhatai* (97%) | - | - | + | - |
| EBL30, 31, 32, 33 | *Bacillus bombysepticus* (97%) | - | - | + | - |
| EBL34, 35,36, 37 | *Bacillus subtilis* (99%) | - | - | + | + |
| EBL38, 39, 40, 41,42 | *Bacillus* sp. S10 (99%) | - | - | + | - |
| EPL1, 2, 3, 4 | *Pseudomonas putida* (98%) | - | + | + | + |
| EPL5, 6, 7 | *Pseudomonas chlororaphis* (99%) | + | - | + | + |
| EPL8, 9, 10, 11 | *Pseudomonas fluorescens* (99%) | - | + | + | + |
| EPL12, 13, 14, 15, 16 | *Pseudomonas cichorii* (98%) | - | - | + | + |

<sup>z</sup> Plant beneficial traits were qualitatively evaluated using the methods reported by Trivedi et al (2011). Antibacterial activity: bacterial ability
to inhibit the growth of *Xanthomonas citri subsp. citri* 306 on nutrient agar (NA) plate; IAA production: production of indole acetic acid; P
- solubilization: phosphate solubilization. “+”: presence of trait; “-”: absence of trait.

### Plant-beneficial traits of enriched bacteria revealed by phenotypic assays and genomics analysis

Next, we investigated whether the bacterial isolates of enriched phyllosphere microbes harbor beneficial properties that potentially enhance plant health and productivity. The results showed that representative isolates of the 14 enriched bacterial species contained a minimum one of four beneficial traits related to antibacterial activity, mineral nutrition acquisition, i.e., P solubilization and siderophore production, and plant growth and development, i.e., production of IAA (**Table 1**). Specifically, the isolates of *Brevibacillus agri* EBL1, *P. chlororaphis* EPL5 and *P. fluorescens* EPL8 harbored three of the four beneficial traits tested. EBL1 and EPL5 had substantial antimicrobial activity against Xcc strain 306 *in vitro* (**Supplementary Figure 6**). Therefore, these three strains were subjected to whole-genome sequencing for further investigations.

General information for the three draft genome sequences is summarized in **Supplementary Table 5**. The mean coverage was 360 for EBL1 in 383 scaffolds, GC content for EBL1 was 53.86% and the total genome size was 5.33 Mb. The annotation identified 5072 CDs and 97 rRNAs and tRNAs in this genome. The mean coverage was 407 for EPL5 in 38 scaffolds and 184 for EPL8 in 1902 scaffolds. EPL5 and EPL8 draft genomes have a GC content of 62.95% and 60.92% and a genome size of 6.78 Mb and 7.09 Mb, respectively. There are 6069 CDs and 70 rRNAs and tRNAs in EPL5 and 5962 CDs and 85 rRNAs and tRNAs in EPL8 draft genomes.

KO-based annotation of the whole genomic sequences revealed that EBL1, EPL5, and EPL8 contain gene clusters involved in P solubilization such as *phoP* encoding a transcriptional regulator and *phoR* coding for a phosphate regulon sensor protein, and nitrogen fixation including *nirA*, *nirB*, and *nirK* (**Fig.3**). EBL1, EPL5, and EPL8 also have gene clusters related to siderophore production and iron acquisition, and production of phytohormones and volatile organic compounds (VOCs).

Genome mining using antiSMASH indicated that EBL1, EPL5, and EPL8 possess differed numbers of gene clusters responsible for biosynthesis of antibiotics and other secondary metabolites, with nine clusters predicted for EBL1, 16 clusters for EPL5, and 13 for EPL8 (**Supplementary Table 6**). EBL1 has three clusters of non-ribosomal peptide synthetase (NRPS) or NRPS-like genes and two polyketide synthase (PKS) genes clusters. One NRPS cluster showed 43% nucleotide sequence similarity to the tyrocidine biosynthetic gene cluster from *Brevibacillus brevis,* a metabolite that has strong antibacterial activity. Another NRPS cluster shared 21% nucleotide sequence similarity with the paenilamicin biosynthetic gene cluster, a polycationic peptide with antibacterial and antifungal activities. The NRPS-like gene cluster had 21% nucleotide sequence similarity with the biosynthetic gene cluster of lichenysin, a cyclic lipopeptide that has antimicrobial activity. EPL5 has four NRPS or NRPS-like genes clusters and one cluster for NRPS-independent, IucA/IucC-like siderophores. Two NRPS clusters exhibited 21% and 34% nucleotide sequence similarity respectively to the pyoverdine biosynthetic gene cluster in *Pseudomonas protegens* Pf-5 (previously called *P. fluorescens* Pf-5) and the NRPS-like cluster had 37% nucleotide sequence similarity to the fragin biosynthetic gene cluster. Notably, EPL5 also carries two gene clusters with 100% similarity to the pyocyanin and pyrrolnitrin biosynthesis clusters. Both pyocyanin and pyrrolnitrin are well-characterized compound produced by many *Pseudomonas* strains, exhibiting antimicrobial activity against various bacteria and fungi. EPL8 has four NRPS or NRPS-like genes clusters and one cluster for NRPS-independent, IucA/IucC-like siderophores. One NRPS cluster had 100% similarity to the biosynthesis cluster for nunapeptin and nunamycin, antifungal secondary metabolites in *P. fluorescens.* Another NRPS cluster had 84% similarity to the biosynthesis cluster for histicorrugatin, a siderophore with antifungal activity produced by *Pseudomonas* spp. Collectively, the results demonstrated that the bacteria *Brevibacillus agri* EBL1, *Pseudomonas chlororaphis* EPL5, and *Pseudomonas fluorescens* EPL8, which were enriched in citrus leaves by microbial manipulation with SynCom1, contain multiple plant-beneficial properties that may promote citrus health and productivity.

### Bacterial inoculation activated citrus defense responses

We further assessed whether single strain (ATY16, PT6, or PT26A) or SynCom1 (ATY16 + PT6 + PT26A) inoculation can activate citrus defense responses. The results showed that PT26A or SynCom1 inoculation triggered differential expression of the defense-related genes in citrus leaves compared to untreated control (**Fig.4**), while ATY16 or PT6 inoculation did not (data not shown). The levels of gene expression induced by SynCom1 were not significantly different from those induced by PT26A. Both PT26A and SynCom1 significantly up-regulated (≥ 2-fold) the expression of genes for the pathogenesis-related proteins 1, 2, and 5 (*PR1, PR2,* and *PR5*), for allene oxide synthase (*AOS*), and plant defensin gene *PDF1.2,* but not the gene for allene oxide cyclase (*AOC*) or basic helix–loop–helix (bHLH) transcription factor (*MYC2*), at 10 to 72 hours post inoculation (**Fig.4**). Peak levels of gene expression were observed at 24 hours post inoculation. Specifically, PT26A and SynCom1 elicited stronger expressions of *PR1, PR2* and *PR5* than *AOS* and *PDF1.2.* PR genes are usually activated through the SA-signaling pathway while the rest of the genes (*AOS*, *AOC*, *MYC2*, and *PDF1*.2) are associated with the JA-signaling pathway. These results suggest that foliar spray of SynCom1 may activate citrus defense response via both the SA and JA pathways.

### Microbiome manipulation reduced citrus canker severity in greenhouse assays

Next, we tested whether pre-inoculation of single strain (i.e., ATY16, PT6, PT26A, EBL1, or EPL5), SynCom1, or SynCom2 (i.e., mixture of ATY16, PT6, PT26A, EBL1, and EPL5) via foliar spray can suppress citrus canker. Here, we have designed SynCom2 by adding *Brevibacillus agri* EBL1, and *Pseudomonas chlororaphis* EPL5 to SynCom1 based on compatibility. In greenhouse assays, the bacterial treatments were applied two days prior to Xcc 306 inoculation, and disease incidence (percentage of plants showing canker symptoms) and severity (average counts of canker lesions per leaf) were estimated at four weeks post Xcc 306 inoculation. The results showed that none of the bacterial treatments reduced the canker disease incidence, as 100% treated plants developed canker lesions on leaves. However, a significant reduction of canker disease severity was observed in the treatments of single strains (ATY16, PT6, or PT26A), SynCom1, and SynCom2, compared with untreated control (*P* < 0.0001; One-way ANOVA) (**Fig. 5A**). Moreover, SynCom1 and SynCom2 had a better efficacy in reducing disease severity than single-strain treatments. There were no significant differences in the efficacy of reducing disease severity between SynCom1 and SynCom2 (**Fig. 5A**). These results indicate that the bacterial inoculations significantly reduced the citrus canker disease severity and SynCom1 and SynCom2 treatments were more effective than any single strains in suppressing citrus canker.

To further evaluate the control effect of different treatments, we quantified Xcc growth in different treatments. At 6 hours (0 DPI), Xcc strain 306 populations in citrus plants pretreated with single strain, SynCom1, SynCom2, or copper were at a similar level of approximately 1-2 × 10^5^ CFU/cm^2^ leaf tissue with those in mock plants (pretreated with water) (**Fig. 5B**). At 7 DPI, the treatments of copper, SynCom1, and SynCom2 had a significantly lower level of Xcc populations at 0.5-1.0 × 10^4^ CFU/ cm^2^ leaf tissue than mock plant at 0.5-1.0 × 10^5^ CFU/cm^2^ leaf tissue, while the treatments with single strains carried similar Xcc populations as mock treatment. At 14 to 28 DPI, Xcc populations in plants pretreated with single strains, SynCom1, SynCom2, or copper were significantly lower than in mock plants. Moreover, copper treatment consistently had the lowest level of Xcc populations, followed by the treatments of SynCom1 and SynCom2, single strains, and mock. At 28 DPI, the differences in population sizes of Xcc between copper, SynCom1, SynCom2 or single strains and mock treatment were three, two, and one order(s) of magnitude, with Xcc strain 306 at about 10^5^, 10^6^, 10^7^, and 10^8^ CFU/cm^2^ leaf tissue in the copper, SynComs, single strains, and mock treated plants, respectively (**Fig. 5B**). Among the single-strain treatments, there were no significant differences in Xcc population sizes from 0 to 28 DPI. Similarly, there were no significant differences in Xcc population size between SynCom1 and SynCom2 from 0 to 28 DPI, except at 14 DPI. Compared with single strains, SynCom1 and SynCom2 significantly reduced Xcc population levels (*P* < 0.0001), and the latter was more effective and comparable to copper (**Fig. 5B**). These results suggest that SynCom1 and SynCom2 performed better than single strains in suppressing Xcc population growth *in planta*. Importantly, Xcc population was similar in the copper and SynCom2 treatments at 21 and 28 DPIs (Fig. 5B).

### Suppression of citrus canker by microbial treatment in the field

Based on the observations in the greenhouse tests, we evaluated the effectiveness of SynCom1 and SynCom2 against citrus canker under field conditions for three years from 2022 to 2025. It is worthy noting that we rotated SynCom1 or SymCom2 with copper hydroxide treatments from the second year, thus these two treatments were only done for two years from 2023-2025. The treatment applications were conducted at a three-week interval from March to October and disease severity of foliar and fruit canker (incidence of canker-infested leaves and fruit) in treated trees was estimated seven months after the first application of treatments. Symptoms of foliar and fruit canker were observed in all experimental trees over the tests. In the 3-year trials, untreated trees had an average 49 to 62% leaves, and 43 to 47% of fruits exhibited canker symptoms. All treatments were effective for reducing incidence of foliar and fruit canker compared to untreated control. SynCom1 and SynCom2 were equally effective for suppressing foliar and fruit canker, with a disease incidence of 24 to 37% and 20 to 23%, respectively (**Table 2**). Moreover, rotation of SynCom1 or SynCom2 with copper hydroxide was more effective than the bacterial mixture alone, resulting in a reduction of foliar and fruit canker incidence to 19 - 21% and 16-19% respectively. The most effective treatment was copper hydroxide applied at a standard rate, decreasing foliar and fruit canker incidence to 13 - 24% and 12-17%, respectively. In other words, copper hydroxide had an average efficacy of 68.3% and 69.2% controlling foliar and fruit canker. SynCom1 and SynCom2 alone showed an equal control efficacy of 49.8% and 51.1% against foliar and fruit canker, and when rotated with copper hydroxide they exhibited an equal control efficacy of 60.1% and 61.5% for foliar and fruit canker, respectively. These results indicate that SynCom1 and SynCom2 consistently suppressed citrus canker under differed environmental conditions (with the occurrence of hurricanes in central Florida in 2022 and 2024), implying that they could be an alternative to copper-based bactericides in citrus canker management.

**Table 2.** Incidence of leaves and fruit with symptoms of citrus canker on 4-, 5-, and 6-year-old ‘OLL-8’ sweet orange trees treated with bacterial synthetic communities (SynComs) and/or copper hydroxide at Lake Alfred, FL in 2022, 2023 and 2024 ^y^.

| Treatment<sup>z</sup> | 2022 |  | 2023 |  | 2024 |  |
| --- | --- | --- | --- | --- | --- | --- |
|  | Canker-infected leaves (%) | Canker-infected fruit (%) | Canker-infected leaves (%) | Canker-infected fruit (%) | Canker-infected leaves (%) | Canker-infected fruit (%) |
| UTC | 62.1 a | 43.5 a | 49.4 a | 42.6 a | 51.7 a | 47.4 a |
| SynCom1 | 35.6 b | 20.9 b | 25.3 b | 20.1 b | 23.5 b | 22.6 b |
| SynCom2 | 37.3 b | 23.1 b | 26.4 b | 19.5 b | 23.9 b | 23.2 b |
| Copper hydroxide | 24.5 c | 16.6 c | 14.7 d | 12.2 d | 13.2 d | 11.6 d |
| SynCom1/Copper hydroxide | NT | NT | 21.1 c | 15.8 c | 19.7 c | 18.6 c |
| SynCom2/Copper hydroxide | NT | NT | 20.6 c | 15.6 c | 19.3 c | 17.9 c |

<sup>y</sup> Leaves and fruits were evaluated in October 2022, November 2023 and November 2024. Values are the means of six replicate plots of five trees. Values within columns followed by different letters are significantly different ( $P < 0.05$ ) based on Tukey’s HSD test ( $\alpha = 0.05$ ). NT = not treated in 2022.
<sup>z</sup> UTC: untreated control (sprayed with water); SynCom1: synthetic community 1 at $1 \times 10^9$ CFU/ml containing *Paenibacillus* sp. ATY16, *Bacillus megaterium* PT6, and *Bacillus subtilis* PT26A in a 1:1:1 mixture (each at $1 \times 10^9$ CFU/ml); SynCom2: synthetic community 2 at $1 \times 10^9$ CFU/ml containing *Paenibacillus* sp. ATY16, *Bacillus megaterium* PT6, *Bacillus subtilis* PT26A, *Brevibacillus parabrevis* EBL1 and *Pseudomonas fluorescens* EPL5 in a 1:1:1:1:1 mixture (each at $1 \times 10^9$ CFU/ml). SynCom1/Copper hydroxide: SynCom1 rotated with copper hydroxide; SynCom2/Copper hydroxide: SynCom2 rotated with copper hydroxide.

## Discussion

In this study, we revealed that the structure of citrus leaf microbiomes can be altered by microbial inoculation with a SynCom, leading to a microbial community with the enrichment of beneficial resident bacteria. In addition, treatments of citrus with SymCom and individual beneficial bacteria showed promising effect in suppression of the citrus bacterial canker disease. SymCom is significantly more efficient in suppressing Xcc than individual beneficial bacteria, which probably results from the complementary and additive mode of action of individual bacteria in pathogen inhibition, recruiting of antagonistic bacteria, activation of plant deference response, and competition with the pathogen for resource (nutrients). Moreover, we demonstrated the effectiveness of biological control of citrus canker by the SynCom under field conditions. Unlike the rhizosphere, phyllosphere is directly exposed to the natural environment and has easy access for agricultural management practices. This work provides novel insights into manipulating the phyllosphere microbiome to manage plant diseases and offers new opportunities to enhance plant health and agricultural sustainability through microbiome engineering.

It is believed that a more complex microbial inoculation (e.g., SynCom) would result in more effective disease suppression, as taxonomic diversity is an important feature of microbiome structure and function (Henry and Bergelson 2025; Hirata et al. 2025; Martins et al. 2023). However, multiple studies suggested that the diversity of microbial inoculum is not directly correlated with the level of disease suppression (Berg and Koskella 2018; Cui et al. 2021). In those investigations, multiple strains belonging to same or different bacterial species were mixed for individual microbial treatments, based on the notion that the diversity of a constructed bacterial community would increase in accordance with the increasing number of strains, without consideration of complementation of their modes of action. In contrast, the present study explored complex inoculations consisting of multiple mechanistically complemented plant beneficial bacterial strains relative to individual strains alone and, the results showed that complex inoculations outperformed single-species inoculations in modulation of leaf microbiome structure and protection against the citrus bacterial canker disease. Our findings, together with previous reports toward biocontrol of plant diseases such as fire blight on pears (Stockwell et al. 2011) and tobacco sudden wilt associated with *Fusarium-Alternaria* (Santhanam et al. 2015; Santhanam et al. 2019), underscored mechanistic complementation of members within a SynCom for successfully manipulating plant microbiome for effective control of plant diseases.

Notably, our results showed that SynCom1 inoculation reshaped the leaf microbiome structure, with the enrichment of resident *Bacillus* spp. and *Pseudomonas* spp. that harbor plant-beneficial traits alike to those of the inoculated strains. One possible reason for these observations could be the changes in localized microbial interactions in the leaves. First, the addition of SynCom1 and subsequent excess proliferation of *Bacillus* spp. might restrain the growth of other leaf endophytic bacteria due to antagonism and/or competition for resources. Indeed, assays for pairwise interactions in the leaves of *Arabidopsis* wild-type Col-0 plants revealed that Firmicutes strains outcompeted Proteobacteria strains at a higher inoculation level (Chen et al 2020). Moreover, consistent with our earlier observations of growth compatibility between citrus endophytic *Bacillus* spp. and *Pseudomonas* spp (Li et al. 2013), we did not detect antibiosis between the enriched *Bacillus* spp. and *Pseudomonas* spp. isolates (data not shown). Consequently, the growth of leaf endophytic bacteria excluding *Pseudomonas* might be limited by *Bacillus* in SynCom1 inoculated leaves. Subsequently, relative to untreated control, more intense bacterial-to-bacterial microbial networks that include complicated connections among specific members of Firmicutes and Proteobacteria could occur in the phyllosphere microbiota of SynCom1 inoculated leaves (**Supplementary Figure 5**). In such a scenario *Bacillus* spp. and *Pseudomonas* spp. could outcompete other endophytic bacteria.

Core members in plant microbial communities play important roles in not only suppressing phytopathogens but also maintaining the antipathogen functions of the whole community in healthy plants. For example, Niu et al. (2017) reported that removing *Enterobacter cloacae* from the constructed bacterial community of seven-species that was effective against maize seedling blight caused by *Fusarium verticillioides* led to the biocontrol community collapse (Niu et al. 2017). Additionally, *Bacillus* spp., including many species that are potentially plant-beneficial, is a core member of the citrus phyllosphere microbiome and citrus plants maybe need to maintain a substantial population size of *Bacillus* spp. to stay healthy (Zhang et al. 2021). Specifically, recent studies suggested that the residents of *Bacillus* spp. in citrus leaves might play a role in suppressing citrus canker disease progression (Huang et al. 2023), and positive associations have been observed between the relative abundances of *Bacillus* spp. within citrus microbiomes and citrus genotypes resistant or tolerant to citrus canker (Xu et al. 2023). Thus, with the concept of ecologically informed microbiome engineering (Delgado-Baquerizo et al. 2025; Henry and Bergelson, 2025), one may speculate that targeted microbiome engineering that seeks to shift the citrus microbial communities to a status containing enriched key members associated with plant health (e.g., *Bacillus* spp.) and maintaining a stable structure may protect the host plant from damage caused by pathogens. Our findings that foliar introduction of native microbes enabled the enrichment of beneficial *Bacillus* spp. and *Pseudomonas* spp. in the phyllosphere microbiome and reduction of citrus canker disease severity are supportive to this notion.

In greenhouse assays, SynCom1 and SynCom2 produced a substantial suppression of the citrus canker symptoms. Particularly, the efficacy of SynCom2 appeared comparable to the copper-based bactericide tested (**Fig.5**), implying that the cooperative interactions of SynCom1 with the enriched resident microbiome members might confer more robust biocontrol effectiveness than SynCom1 alone. However, this observation was not reproduced in the field tests. Some possible reasons could be the differences in frequency of pathogen inoculation (i.e., single inoculation of pathogen in greenhouse versus repeated inoculations by windstorms in the field after treatment application) and differences in local environmental conditions that might impact the plant-microbiome-pathogen interactions. Similarly, previous studies have shown that many biocontrol bacteria had inconsistent effectiveness under field conditions, although they were determined with excellent biocontrol potential under *in vitro* or *in planta* conditions (Johnson and Stockwell 1998; Sundin et al. 2009). Therefore, further field tests are needed.

The findings in the present study highlight that microbiome manipulation-based strategy could be a viable alternative to copper-based bactericides for control of citrus canker. All manipulations employed in this study produced substantially decrease in citrus canker severity in the field, although none could achieve the same disease suppression as the copper-based bactericide (**Table 2**). Furthermore, when rotated with copper hydroxide, both SynCom1 and SynCom2 had a control efficacy (60%) approaching that (70%) achieved by copper hydroxide applications. This meant that reducing copper hydroxide usage to half of the standard season-long applications by alteration with the bacterial consortium was still highly effective in controlling citrus canker under field conditions. This deserves further investigations for a better efficacy as it will reduce the potential adverse impact of copper on the environment and thus promote agricultural sustainability.

One limitation of this study is that the field trials were conducted in one citrus grove with a single variety. It remains to be determined whether the biocontrol efficacy of the bacterial consortium against citrus canker varies on different citrus varieties or at differed locations. Nevertheless, the field trials in this study were implemented for three consecutive years and in each year, there were different climate conditions. Thus, the results and findings are scientifically sound and meaningful. Foliar plant pathogens pose increasing threats for agricultural crop production, especially fruit and vegetables worldwide due to frequent warm and moisture conditions associated with climate change (Miller et al., 2024). Particularly, some *Xanthomonas* pathogens, with the tropospheric ozone (O_3_) stress, could produce more severe diseases on a resistant cultivar of pepper than under the conditions without O_3_ stress, by altering interactions among microbial community members in the phyllosphere and overall community functions (Bhandari et al., 2023). Therefore, understanding the approaches and mechanisms that drive the assembly of pathogen-suppressive phyllosphere microbiomes in agricultural cropping systems is crucial to microbiome engineering toward effective disease management for sustainable agricultural production. While several studies have claimed the significance of manipulating microbial communities in the development of suppressive capabilities in the phyllosphere, the current work highlights that manipulation-conferred, targeted impacts on bacterial community composition and specific interactions within the leaf endophytic microbial community could be important players in achieving disease suppression and maintaining plant health. In conclusion, our findings provide novel insights and valuable clues for microbiome engineering to protect agricultural crops from pathogen attack in the phyllosphere.

## AUTHOR CONTRIBUTIONS

NW and JL conceptualized and designed the experiments. JL, JX, DL, ZP, WW, MACG, YF, JLD, YD, MZ CO, and NW performed the experiments. JL, JX, ZP, and NW conducted data analysis. JL, JX and NW wrote the manuscript.

## Acknowledgments

We appreciate the collaborative grower for allowing us to perform the field experiments in one of his citrus groves at Lake Alfred, Florida.

## Funding

This work was supported by the US Department of Agriculture-National Institute of Food and Agriculture (USDA-NIFA) Agriculture and Food Research Initiative Program (grant number 2021-67013-34588 to N.W.) and Hatch Project FLA-CRC-005979 (N. Wang). The funders had no role in experiment design, data collection and analysis, decision to publish, or preparation of the manuscript.

## Supplementary Materials

**Supplementary Figure 1.**
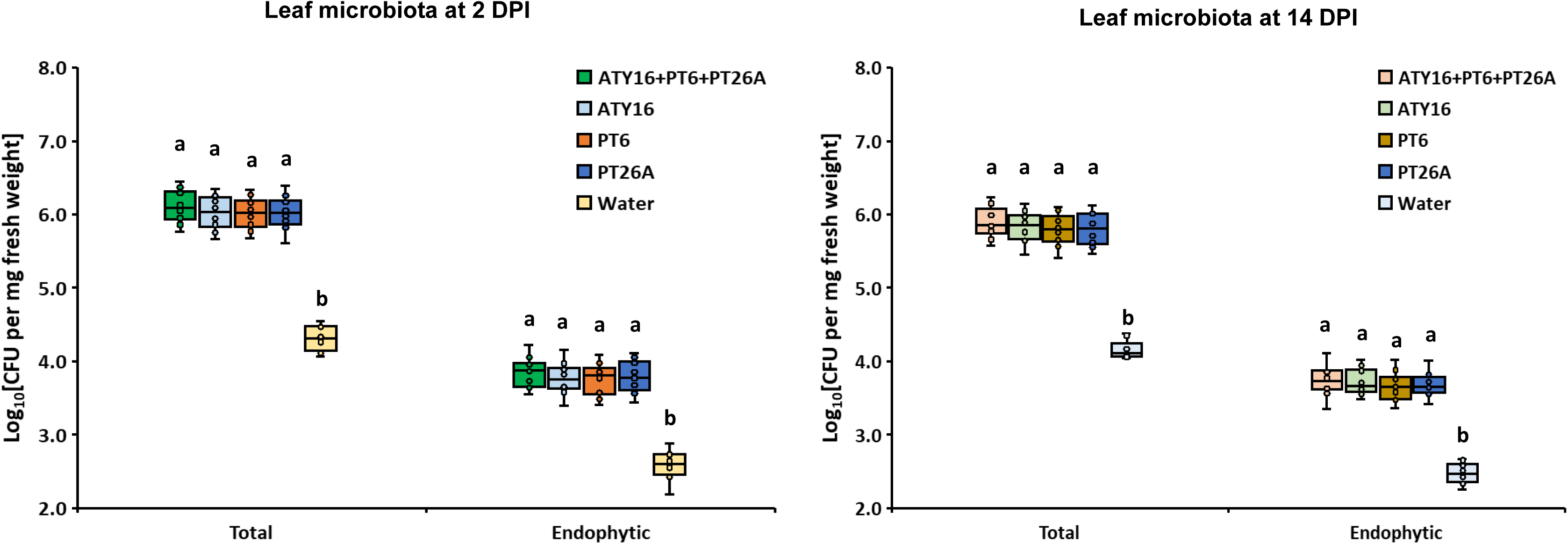
Population sizes of total and endophytic leaf microbiota in citrus plants received different bacterial inoculations in greenhouse. Data shown are combination of results of two independent experiments. Different lowercase letters indicate significant differences in total or endophytic population between the groups (SynCom1, ATY16, PT6, PT26A, and water) (*P* < 0.05, One-way ANOVA with Tukey’s HSD test at α = 0.05). *n* = 4 biological replicates. In box plots, the center line denotes the median, box edges represent the 75^th^ and 25^th^ percentiles, and whiskers show 1.5× the interquartile range.

**Supplementary Figure 2.**
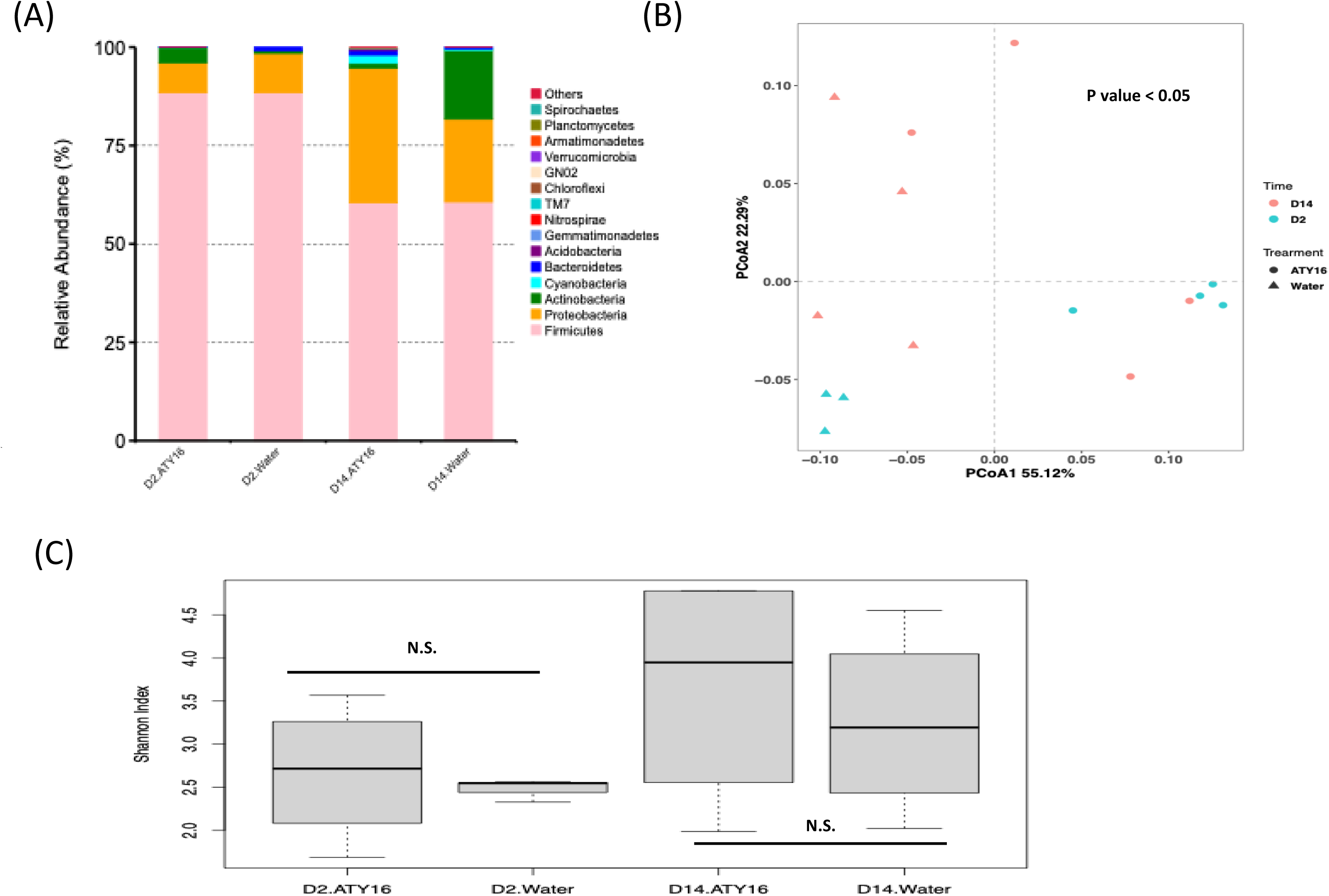
Endophytic leaf microbiomes in strain ATY16 sprayed and water treated control plants. (A) The relative abundance of bacteria at the phylum level in citrus plants sprayed with strain ATY16 or water across time. (B) Principal coordinate analysis (PCoA) based on the weighted UniFrac distance in strain ATY16 treatment and control samples from each timepoint. *P* value was generated by permutational multivariate analysis of variance (PERMANOVA); (C) Shannon indexes (alpha diversity) calculated at the genus level based on 16S rRNA gene-sequence profiles of endophytic leaf bacteria in citrus plants sprayed with strain ATY16 or water. N.S.: no significant differences (*P* > 0.05; Tukey’s HSD test at α = 0.05) *n* = 3 biological replicates (water treatment at 2 days post inoculation (DPI)) and *n* = 4 biological replicates (ATY16 treatment at 2 and 14 DPI and water treatment at 14 DPI). D2: 2 DPI. D14: 14 DPI. Water: water treatment as untreated control.

**Supplementary Figure 3.**
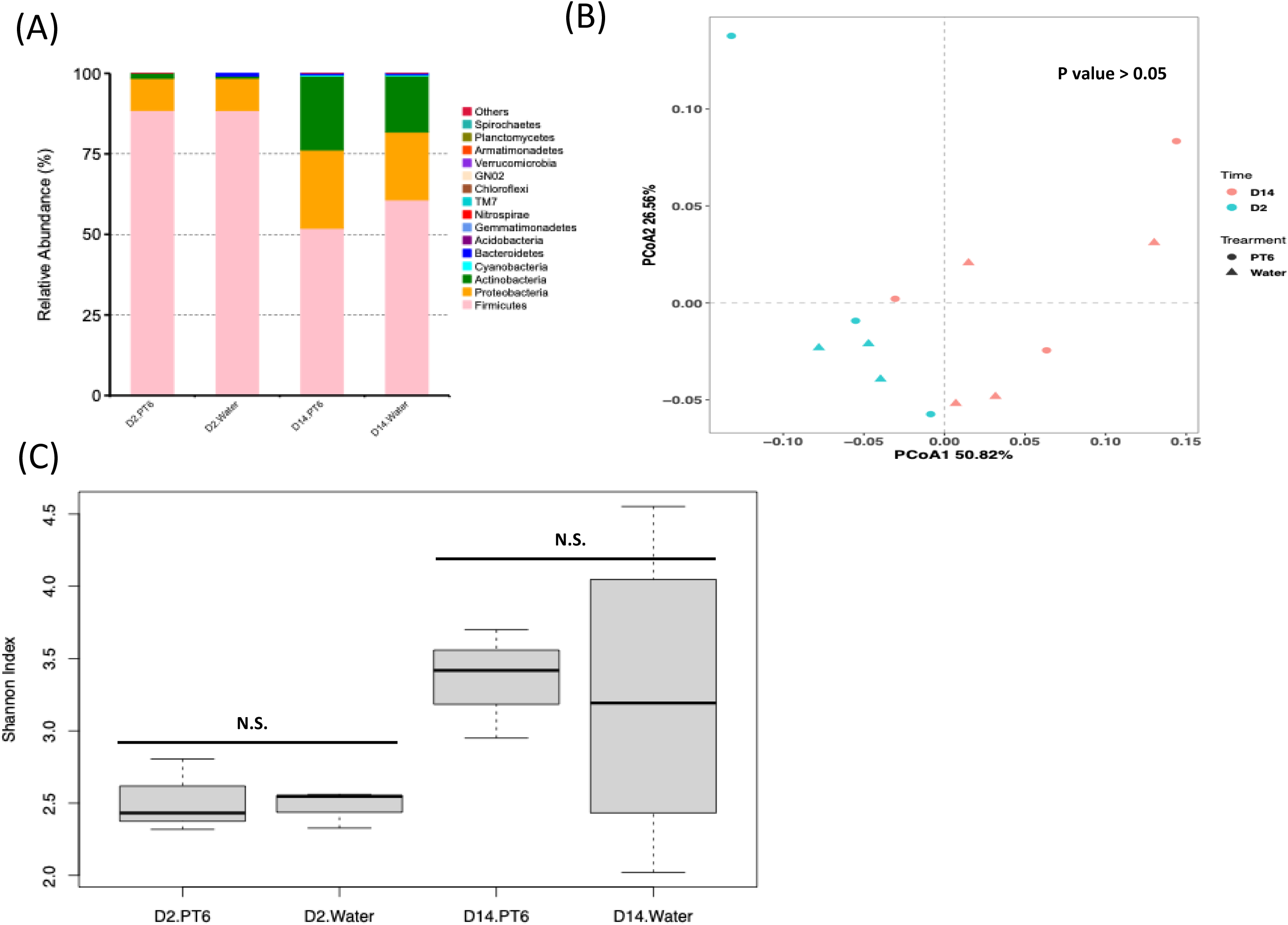
Endophytic leaf microbiomes in strain PT6 sprayed and water treated control plants. (A) The relative abundance of bacteria at the phylum level in citrus plants sprayed with strain PT6 or water across time. (B) Principal coordinate analysis (PCoA) based on the weighted UniFrac distance in strain PT6 treatment and control samples from each timepoint. P-value was generated by permutational multivariate analysis of variance (PERMANOVA); (C) Shannon indexes (alpha diversity) calculated at the genus level based on 16S rRNA gene-sequence profiles of endophytic leaf bacteria in citrus plants sprayed with strain PT6 or water. N.S.: no significant differences (*P* > 0.05; Tukey’s HSD test at α = 0.05) *n* = 3 biological replicates (water treatment at 2 days post inoculation (DPI)) and *n* = 4 biological replicates (PT6 treatment at 2 and 14 DPI and water treatment at 14 DPI). D2: 2 DPI. D14: 14 DPI. Water: water treatment as untreated control.

**Supplementary Figure 4.**
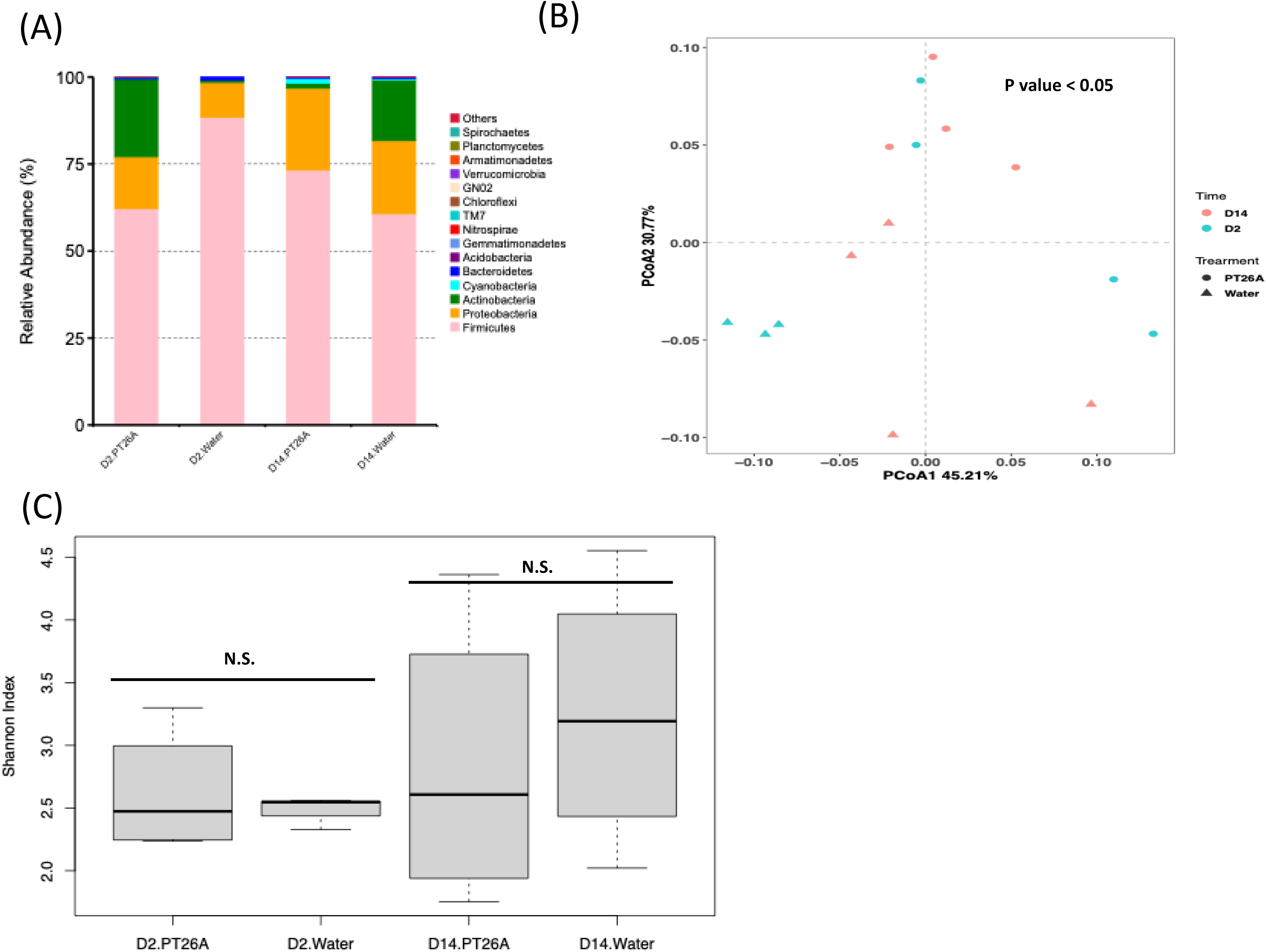
Endophytic leaf microbiomes in strain PT26A sprayed and water treated control plants. (A) The relative abundance of bacteria at the phylum level in citrus plants sprayed with strain PT26A or water across time. (B) Principal coordinate analysis (PCoA) based on the weighted UniFrac distance in strain PT26A treatment and control samples from each timepoint. P-value was generated by permutational multivariate analysis of variance (PERMANOVA); (C) Shannon indexes (alpha diversity) calculated at the genus level based on 16S rRNA gene-sequence profiles of endophytic leaf bacteria in citrus plants sprayed with strain PT26A or water. N.S.: no significant differences (*P* > 0.05; Tukey’s HSD test at α = 0.05) n = 3 biological replicates (water treatment at 2 days post inoculation (DPI)) and *n* = 4 biological replicates (PT26A treatment at 2 and 14 DPI and water treatment at 14 DPI). D2: 2 DPI. D14: 14 DPI. Water: water treatment as untreated control.

**Supplementary Figure 5.**
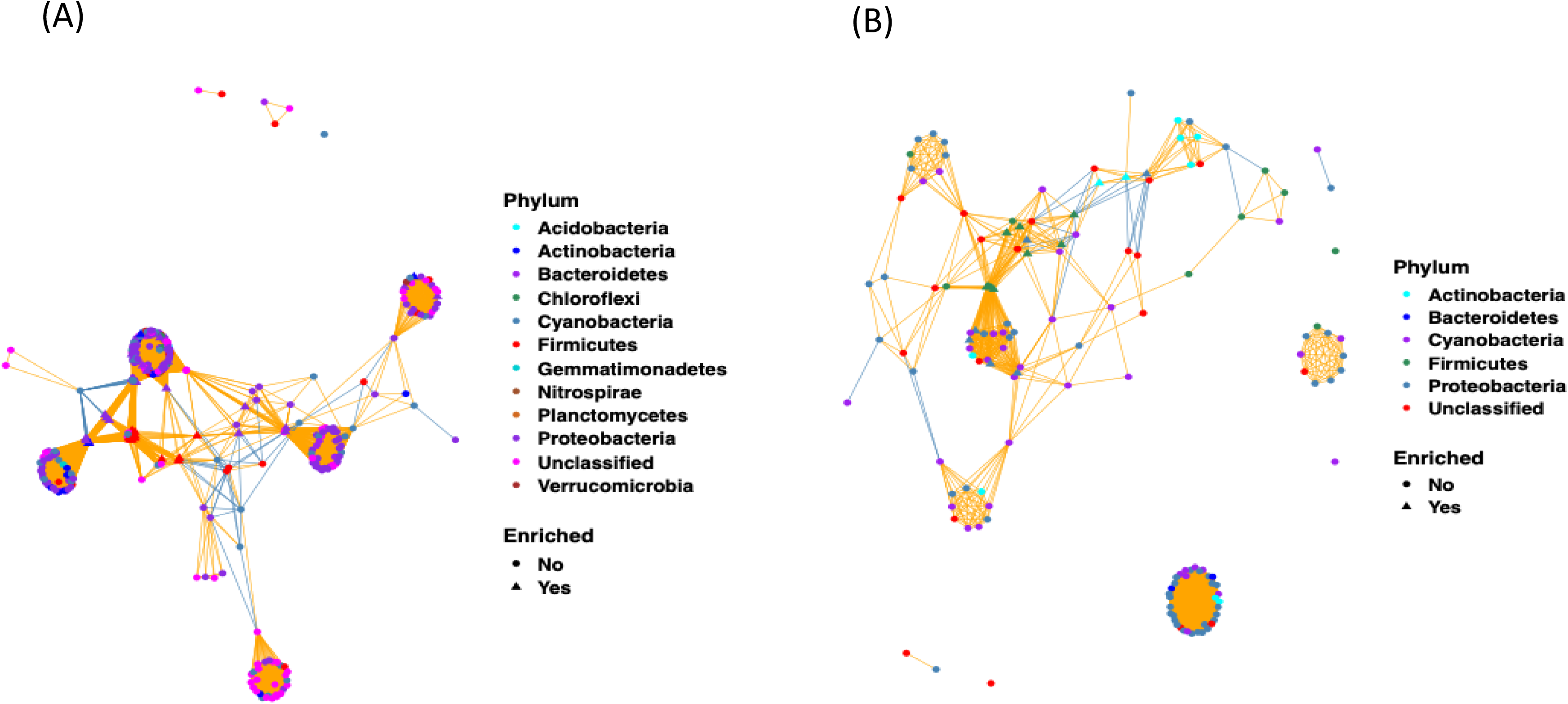
Bacterial co-occurrence networks in the endophytic leaf microbiomes in the bacterial mixture SynCom1 sprayed and water treated citrus plants at 2 (A) and 14 (B) days post inoculation. The color of edge lines indicates the correlation among ASVs: positive correlation (orange) or negative correlation (steel blue). Enriched: the ASVs showed significantly higher abundance in SynCom1 treated samples than control samples.

**Supplementary Figure 6.**
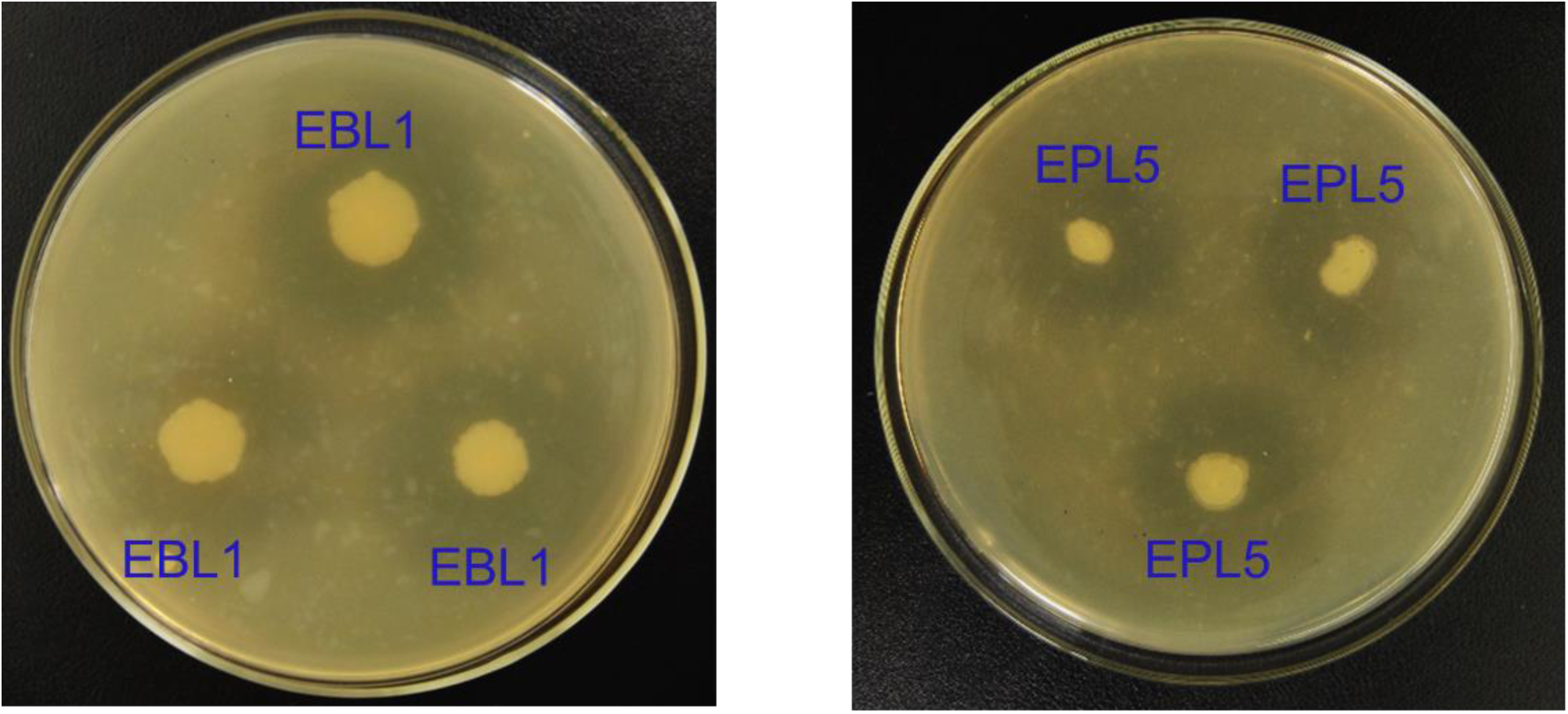
Antibacterial activity of *Brevibacillus parabrevis* EBL1 and *Pseudomonas chlororaphis* EPL5 against *Xanthomonas citri*. ssp. *citri* strain 306 on nutrient agar plates. Both EBL1 and EPL5 contain three replicates in individual plates and inhibition zones occur around each replicate.

**Supplementary Table 1.** Citrus endophytic bacteria used for microbiome manipulation in this study

**Supplementary Table 2.** Primers used for expression analysis of plant defense genes in citrus

**Supplementary Table 3.** Bacteria with different relative abundance between the endophytic leaf microbiomes under the bacterial mixture (SynCom1) and water control treatments at 2 days post inoculation

**Supplementary Table 4.** Bacteria with different relative abundance between the endophytic leaf microbiomes under the bacterial mixture (SynCom1) and water control treatments at 14 days post inoculation

**Supplementary Table 5.** General features of the three draft genomes of sequenced endophytic bacteria

**Supplementary Table 6** Secondary metabolites biosynthesis gene clusters in three bacterial isolates predicted by antiSMASH

## Notes

### Competing Interest Statement

The authors have declared no competing interest.

